# Computational Inference of Metabolic Programs: A Case Study Analyzing the Effect of BRCA1 Loss

**DOI:** 10.1101/2025.11.18.689039

**Authors:** Maria Masid, Ioanna A. Rota, David Barras, Flavia De Carlo, Pierpaolo Ginefra, Mathieu Desbuisson, Yaquelin Ortiz-Miranda, Dmitriy Zamarin, Sohrab P. Shah, Nicola Vannini, George Coukos, Denarda Dangaj-Laniti, Vassily Hatzimanikatis

## Abstract

Metabolic reprogramming is a hallmark of cancer, yet how oncogenic drivers shape tumor metabolism across disease progression remains incompletely understood. In this study, we present iMSEA (*in silico* Metabolic State and Enrichment Analysis), a computational framework that infers flux-based metabolic states from omics profiles. Applying iMSEA to isogenic *BRCA1*-mutant and *BRCA1*-wild-type ovarian cancer cells, we identified a shift toward glycolysis, nucleotide biosynthesis, and redox imbalance, coupled with impaired oxidative phosphorylation. These predictions were validated with metabolomics, Seahorse, and SCENITH assays, demonstrating the accuracy of our approach. Extending the analysis to homologous recombination deficient patient tumors at single-cell resolution, we found that BRCA1-deficient cancers display heightened metabolic activity and site-specific adaptations, including altered central carbon fluxes, mitochondrial function, nucleotide biosynthesis, and lipid metabolism. By linking transcriptional programs to functional metabolic states, iMSEA reveals hidden metabolic liabilities in BRCA1-deficient ovarian cancer and provides a broadly applicable strategy for dissecting metabolic heterogeneity and therapeutic vulnerabilities in cancer.

## INTRODUCTION

Metabolism lies at the heart of tumor behavior, shaping how cancer cells grow, survive, adapt, and respond to therapy. Far from being a passive reflection of genetic changes, metabolic reprogramming is now recognized as a hallmark of cancer, enabling malignant transformation and progression through altered energy production, biosynthesis, and redox balance^1,2^. Understanding these metabolic shifts is fundamental to decoding cancer biology and identifying therapeutic vulnerabilities.

The complexity of cellular metabolism, comprising thousands of interconnected reactions regulated across multiple layers of gene expression, protein abundance, and metabolite availability, demands a systems-level approach to analyze and understand metabolic behavior. Computational systems biology provides a powerful platform for studying metabolism and unraveling the interplay between genetic mutations and metabolic adaptations. Central to this approach are genome-scale metabolic models (GEMs), which mathematically represent the full repertoire of biochemical reactions in a cell and enable the study of cell proliferation, metabolic shifts, and the mechanistic connections between genotype and phenotype^3,4^. Human GEMs such as Recon3D^5^ and Human1^6^ encompass all known metabolic biochemistry in human cells. To study specific contexts, omics data, including transcriptomics, proteomics, and metabolomics, are integrated into these models to build context-specific networks that more accurately reflect the physiological state of interest^7,8^.

Several computational methods have been developed to integrate and interpret omics data using GEMs. Algorithms such as GIMME^9^, iMAT^10^, and MBA^11^ use gene expression thresholds and reaction activity scores to prune the global network into subnetworks that reflect molecular profiles while maintaining metabolic feasibility. More recent approaches, including INIT^12^, tINIT^13^, and REMI^14^ incorporate transcriptomic, proteomic, and metabolomic data to improve tissue-specific model fidelity. Alternative strategies such as mCADRE^15^ and FASTCORE^16^ construct consistent models by expanding from high-confidence reaction cores, offering robustness and scalability in large-scale reconstructions. Focusing on retaining essential metabolic functions during model simplification, redHUMAN^17^ introduced a task-based reduction framework that generates context-relevant, compact metabolic networks. This approach preserves essential biochemical capabilities, avoids over-reduction, and enhances the biological interpretability of the resulting models.

Despite these advances, defining reliable assumptions for integrative analysis and inferring functional metabolic states remains a major challenge. Translating expression data into functional metabolic activity is nontrivial due to uncertainties in gene-reaction mappings and post-transcriptional regulation^18^. Moreover, current approaches often lack systematic frameworks for interpreting predicted metabolic fluxes and rarely incorporate experimental validation to link model outputs with measurable phenotypes^19^. As a result, reliable inference of metabolic states and their connection to cellular behavior remains an open problem in the field.

Bridging the gap between omics data and functional metabolic interpretation requires computational strategies that move beyond static snapshots of gene expression. Ideally, such approaches would infer context-specific metabolic states from high-throughput datasets, while offering interpretable links to phenotypic traits such as proliferation, metabolic flexibility, or nutrient dependence. Crucially, this demands frameworks that integrate multiple omics layers, account for biochemical constraints, and enable systematic hypothesis generation about metabolic reprogramming.

A particularly relevant context in which such an approach is needed is the metabolic rewiring associated with foundational oncogenic mutations. Among these, loss of BRCA1 is a defining feature of high-grade serous ovarian cancer (HGSOC), the most common and lethal subtype of ovarian cancer, characterized by its aggressive nature and poor prognosis^20^. *BRCA1* mutations are prevalent in HGSOC^21,22^, impairing homologous recombination repair of DNA double-strand breaks and promoting genomic instability^23,24^. Beyond defects in DNA repair, BRCA1 loss triggers transcriptional reprogramming, cell-intrinsic inflammation, immune resistance, and tumor progression, profoundly shaping the tumor phenotype^25,26^.

Emerging evidence implicates *BRCA1* as a regulator of endocrine and metabolic pathways, expanding its role beyond genome maintenance^27^. Consequently, mutations in *BRCA1* have been linked to metabolic reprogramming^28,29^, including increased lipid synthesis^30,31^, context-dependent shifts in mitochondrial metabolism^32,33^, and resistance to oxidative stress^34^ to support tumor progression. These metabolic alterations directly impact HGSOC growth, survival, and response to treatment^30^, presenting both significant challenges and potential opportunities for targeted therapies^32,33^. While there is extensive knowledge about *BRCA1* in cancer biology^35^, metabolic rewiring caused by BRCA1 loss at a systems level remains poorly understood. Recent studies have emphasized the role of metabolic heterogeneity in shaping BRCA1-deficient tumor behavior^36,37^, underscoring the need for a comprehensive approach to characterize these alterations in diverse tumor contexts.

In this study, we introduce iMSEA (*in silico* Metabolic State and Enrichment Analysis), a computational systems biology framework designed to identify cellular metabolic states consistent with a given set of omics data and assess cellular phenotypes through enrichment analysis. Using iMSEA, we first investigated the metabolic reprogramming occurring in *BRCA1* isogenic ovarian cancer cell lines. To do so, we integrated gene expression and proteomics data into an ovarian cancer metabolic network, and generated context-specific models representing the metabolism of *BRCA1*-mutant and *BRCA1* wild-type ovarian cancer cells. Using these models, we inferred *in silico* fluxomics that aligned with the transcriptomics and proteomics profiles and identified the metabolic states that emerge in BRCA1 deficient cells. Upon validating the iMSEA predictions by metabolic assays in ovarian cancer cell lines, we applied our framework to investigate metabolic heterogeneity in patient-derived samples. By comparing *BRCA1* mutant-like and wild-type-like tumors across primary ovarian sites and metastatic peritoneal niches, we identified heterogeneous metabolic states and site-specific adaptations. Our analysis demonstrates the utility of iMSEA in uncovering metabolic vulnerabilities and provides a basis for exploring therapeutic interventions tailored to the metabolic profiles of tumor subpopulations.

## RESULTS

### BRCA1 loss provides a suitable model to investigate large-scale metabolic reprogramming

To investigate the impact of BRCA1 loss in the metabolism of HGSOCs, we analyzed the *BRCA1*-null UWB1.289 (*BRCA1*^MUT^) human ovarian cancer cell line, alongside its isogenic counterpart with restored *BRCA1* wild-type (*BRCA1*^WT^) expression^38^ (**Fig. S1a**). The UWB1.289 cell line carries a germline *BRCA1* mutation in exon 11 and a deletion of the wild-type allele, resulting in a truncated nonfunctional BRCA1 protein^38^. In contrast, the UWB1.289+BRCA1 cell line has been engineered to express a full-length, functional *BRCA1* gene, restoring normal protein function and DNA repair capabilities (**Fig. S1b**). Moreover, both cell lines have a somatic *TP53* mutation.

Although no significant differences were observed in growth rates between the *BRCA1*^WT^ (doubling time = 29h) and *BRCA1*^MUT^ (doubling time = 34h) cell lines (**Fig. S1c-d**), gene set variation analysis (GSVA) of hallmark gene signatures revealed multiple alterations in several pathways including activation of angiogenesis and inflammation by BRCA1 loss, in agreement with previous findings^25^. Interestingly, loss of BRCA1 also affected several metabolic pathways. Specifically, *BRCA1*^MUT^ cells exhibited downregulation of pathways related to cholesterol metabolism, along with key metabolic regulators such as mTORC1 and NOTCH (**Fig. S1e**). Complementing these findings, differential gene expression analysis identified significant changes in metabolic genes, including the upregulation of the glycolytic gene lactate dehydrogenase (*LDHC*) and genes from nucleotide (*AMPD3, NT5C1B*) and lipid metabolism (*ACSL5, DGKG, CHPT1*), as well as the downregulation of mitochondrial function genes (*NDUFA2, MT-ND6, UQCRQ*), the fatty acid oxidation rate limiting gene carnitine palmitoyltransferase (*CPT1A*) and oxidative stress-related genes (*SOD3, ALDH4A1, GRHPR*) (**Fig. S1f**). Thus, while cell proliferation *in vitro* remained comparable, loss of BRCA1 produced sufficient gene alterations heralding in principle metabolic reprogramming of these cells, thereby providing a suitable model to test our novel method.

### Computational modeling uncovers metabolic cell states at a systems level

To infer how a single foundational oncogenic alteration, such as BRCA1 loss, globally impacts cellular metabolism, we developed iMSEA (*in silico* Metabolic State and Enrichment Analysis), a computational framework that integrates transcriptional data into genome-scale metabolic models (GEMs) to infer flux-based metabolic states (**Fig. 1a**). Unlike traditional enrichment analyses, iMSEA mechanistically translates gene-expression programs into reaction-level fluxes, providing a systems-level view of metabolic reprogramming.

**Figure 1.**
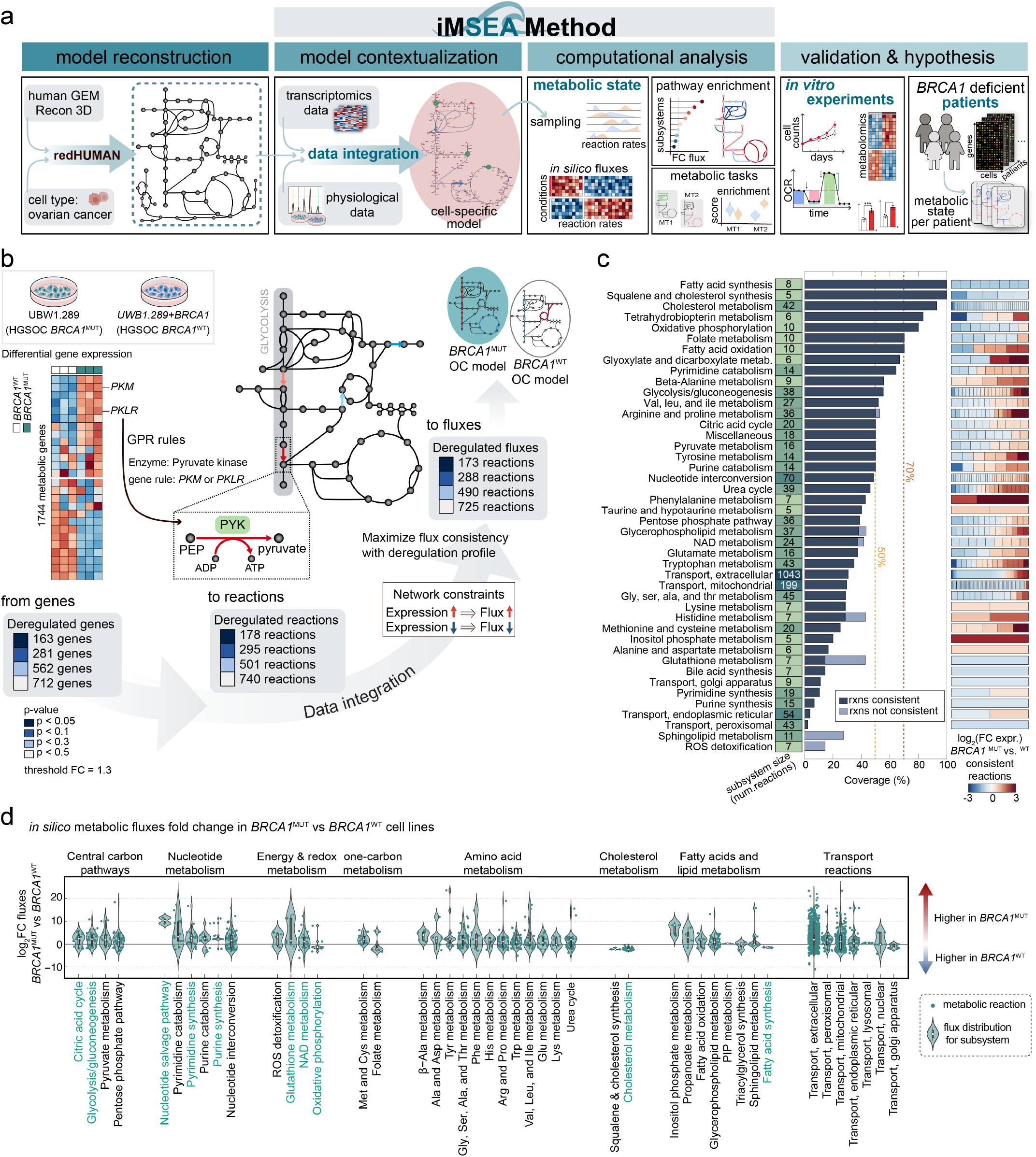
Data integration into metabolic network and inference of metabolic state. **a**, Overview of the workflow to reconstruct context-specific metabolic models and infer metabolic states using iMSEA. **b**, Gene expression data integration into the ovarian cancer (OC) metabolic network with the iMSEA method allowing to infer metabolic fluxes consistent with the gene deregulation profile. Genes are mapped to reactions using the gene-protein-reaction (GPR) rules available in the Recon 3D metabolic network. Reaction rates (fluxes) are constrained based on the corresponding gene expression, assuming a change in expression (up or down) may lead to a change in enzyme availability and therefore a change in flux (up or down). **c**, Metabolic subsystems size, percentage of reactions consistent with gene expression data, and fold change expression of consistent reactions, lower (blue) or higher (red) in *BRCA1*^MUT^ vs. *BRCA1*^WT^ UWB1.289 OC cells. **d**, Reaction rates computationally inferred with iMSEA for all the reactions in the OC metabolic network, classified by subsystem.

iMSEA requires as input a metabolic model that captures the metabolic capabilities of the biological system under study. This model can be generated using established reconstruction or reduction approaches^13,15,16^. In our study, we used the redHUMAN method^17^ to extract a task-relevant subnetwork from the global human metabolic reconstruction Recon3D^5^, focusing on core pathways and physiological functions relevant to ovarian cancer metabolism (**Fig. 1a** and **Methods**). This approach ensured that the resulting model preserved essential metabolic functions, including nutrient uptake, biomass synthesis, and energy production, while reducing complexity to focus on biologically relevant features of ovarian cancer metabolism and improve model interpretability.

We then applied iMSEA to contextualize the ovarian cancer metabolic network using experimental data from *BRCA1*^MUT^ and *BRCA1*^WT^ cells, including RNA-seq profiles, cell replication rates, and nutrient availability (based on culture medium composition). iMSEA addresses a fundamental systems biology challenge: gene-expression changes alone do not reliably predict metabolic activity, since fluxes are shaped by enzyme availability, regulatory constraints, and metabolite supply. By embedding transcriptional data within the stoichiometric and biochemical structure of GEMs, iMSEA generates context-specific models that capture both experimental observations and underlying metabolic mechanisms (**Fig. 1a**).

iMSEA first maps differentially expressed genes (DEGs) to metabolic reactions using gene-protein-reaction (GPR) rules, which describe how genes encode enzymes and how those enzymes catalyze reactions within the network (**Methods**). In the comparison between *BRCA1*^MUT^ and *BRCA1*^WT^ cells, hundreds of metabolic reactions were linked to DEGs, identifying candidate reactions that operate differentially between the two cell states (**Fig. 1b**). A common approach is to interpret DEGs in a point-wise fashion, assuming that if a gene is upregulated its associated reaction must carry higher flux, and if it is downregulated the flux must be lower. For example, *PKM*, which is upregulated in *BRCA1*^MUT^ cells, constrains the pyruvate kinase (PYK) reaction to carry proportionally higher flux in *BRCA1*^MUT^ relative to *BRCA1*^WT^ (**Fig. 1b**). However, across a large interconnected network, many of these point-wise hypotheses cannot all hold true simultaneously, because the system must still satisfy stoichiometric and thermodynamic balances. iMSEA, through its REMI component^14^, resolves this by constructing parallel networks for *BRCA1*^WT^ and *BRCA1*^MUT^ cells and identifying the largest set of flux changes that can be imposed consistently with DEG fold changes while keeping the overall network feasible (**Fig. S2a**). This is formulated as a mixed-integer linear programming (MILP) problem, where binary variables act as switches to maximize the number of gene-flux fold similarities that can be satisfied without forcing impossible changes that would violate the mass balances of metabolites in the network. Through this design, iMSEA quantitatively reconciles gene-level changes with reaction-level flux potential, yielding context-specific models that remain both mechanistically rigorous and biologically interpretable.

A key feature of iMSEA is that gene-expression constraints are integrated sequentially, prioritizing reactions linked to genes showing the strongest statistical significance (**Methods**). The user can define adjusted p-value thresholds to stratify DEGs (e.g., p < 0.05, 0.1, 0.3, and 0.5 in this study), allowing the network to be progressively refined. While strict thresholds (p < 0.05) ensure statistical rigor, we also incorporated genes with strong fold changes at more relaxed p-values (up to 0.5) to capture biologically meaningful trends that might otherwise be masked by variability. This flexibility is particularly relevant in metabolic modeling, where modest or heterogeneous expression changes can still drive important shifts in flux^18^, and predictions can then be evaluated using targeted metabolic assays for validation. Finally, the same mathematical framework extends naturally to comparisons across more than two conditions: iMSEA constructs a condition-specific metabolic network for each context and links them through pairwise comparisons of reaction fluxes (**Fig. S2b**). This design enables detection of both shared and condition-specific metabolic shifts across multiple phenotypic states, broadening the experimental scenarios that iMSEA can address.

### iMSEA reveals metabolic cell states associated with *BRCA1* deficiency

Through this process, we generated two flux-constrained models, one for *BRCA1*^WT^ and one for *BRCA1*^MUT^, representing the inferred metabolic state of the cells (**Fig. 1b**). In the *BRCA1*^MUT^ model, 178 reactions associated with 163 genes were significantly deregulated (p < 0.05). Of these, 173 reactions spanning 31 metabolic subsystems (**Fig. S2c**) were inferred to carry flux values consistent with the direction and magnitude of their gene expression changes. In total, iMSEA constrained the activity of 725 reactions (out of 740 differentially expressed reaction-gene pairs, corresponding to 712 genes), yielding two distinct context-specific metabolic models for *BRCA1*^WT^ and *BRCA1*^MUT^ cells *in vitro* (**Fig. 1b**).

Analysis of the metabolic subsystems associated with the 725 reaction rates inferred to be operating differentially, revealed that the loss of BRCA1 affected key metabolic pathways. Most reactions involved in fatty acid synthesis and cholesterol metabolism were downregulated, with consistently lower fluxes in *BRCA1*^MUT^ compared to *BRCA1*^WT^ cells (**Fig. 1c**). Oxidative phosphorylation (OXPHOS) was also strongly affected, with more than 70% of its reactions aligned with the observed gene downregulation. Conversely, fatty acid oxidation and pyrimidine catabolism were upregulated, with 70% of their reactions carrying higher fluxes in *BRCA1*^MUT^ cells (**Fig. 1c**). In contrast, central carbon pathways, including glycolysis, the pentose phosphate pathway (PPP), and the citric acid cycle (TCA), showed only partial consistency, with 50% of their reaction rates reflecting a combination of up and downregulated genes (**Fig. 1c**). This partial consistency, arising from limited gene coverage and heterogeneous expression trends, highlights the difficulty of drawing definitive conclusions about the activity of these pathways.

Even after integrating DEGs with network balance, multiple feasible flux states remain possible. This reflects a fundamental property of metabolic networks: many different distributions of reaction rates can satisfy the same constraints. To avoid over-interpreting a single mathematical solution, iMSEA explores this space through a final step based on flux sampling (**Methods**). A key technical challenge is that flux sampling methods generally require the solution space to be convex. In a convex space, if two different flux profiles (that is, two different patterns of reaction rates across the metabolic network) are feasible, then any weighted combination of them is also feasible. By contrast, adding gene-based constraints makes the solution space irregular, or non-convex, meaning that some of these combinations are no longer valid. To manage this, iMSEA partitions the flux space into smaller convex regions and applies Markov Chain Monte Carlo sampling using the Artificial Centering Hit-and-Run method within each region. This approach is particularly well-suited for exploring high-dimensional flux spaces under metabolic constraints^39^. Sampling across all regions ensures coverage of the full range of feasible solutions without bias. In total, this procedure generates thousands of flux configurations across regions that are consistent with stoichiometric, environmental, thermodynamic, and gene expression-derived constraints (**Fig. S2d**). From these sampled distributions, we can assess for each pathway whether fluxes are consistently higher or lower in *BRCA1*^MUT^ compared to *BRCA1*^WT^ cells, thereby identifying the subsystems most likely to be deregulated.

As a result, we obtained flux distributions for each metabolic subsystem in the BRCA1 case study (**Fig. 1d**). This analysis confirmed strong downregulation of oxidative phosphorylation, fatty acid synthesis, and cholesterol metabolism, with 80–100% of reactions in these pathways carrying lower flux in *BRCA1*^MUT^ cells, consistent with gene expression trends. By contrast, changes in central carbon metabolism, nucleotide metabolism, glutathione metabolism, and NAD metabolism showed predominantly higher activity in *BRCA1*^MUT^ cells. Notably, the deregulation in these pathways could not be confidently interpreted from gene expression data alone, due to incomplete coverage and mixed up- and downregulated genes. By integrating expression changes within the stoichiometric network, iMSEA provided flux estimates for all reactions, allowing us to resolve these ambiguities. As a result, we inferred that more than 70% of glycolytic and tricarboxylic acid cycle reactions, over 88% of nucleotide synthesis reactions, more than 85% of glutathione metabolism reactions, and approximately 80% of NAD metabolism reactions carried higher flux in *BRCA1*^MUT^ cells. Together, these findings suggest that BRCA1 loss leads to mitochondrial dysfunction coupled with increased anabolic activity, elevated oxidative stress, and higher demand for redox buffering and DNA repair capacity.

### BRCA1 deficiency induces energy metabolism rewiring and redox imbalance in ovarian cancer cell lines

An advantage of iMSEA is its ability to generate detailed metabolic profiles, enabling us to examine not just pathways but also individual reactions within the network (**Fig. 2a**). Our computational model predicted higher flux through both glycolysis and the tricarboxylic acid (TCA) cycle in *BRCA1*^MUT^ cells, consistent with prior studies showing that BRCA1 loss enhances glycolytic activity in cancer cells^36,40^. In our system, this was supported by the upregulation of several glycolytic genes, including hexokinase (*HKDC1*), phosphofructokinase (*PFKM*), enolase (*ENO3*), and acylphosphatase (*ACYP2*), which contributed to the inferred changes in metabolic flux (**Fig. 2b**). To test these predictions experimentally, we first performed intracellular and extracellular metabolomics under basal conditions (**Methods**). While glycolytic intermediates showed little change between *BRCA1*^MUT^ and *BRCA1*^WT^ cells, several TCA intermediates (succinate, malate, and α-ketoglutarate) were elevated in *BRCA1*^MUT^ cells, whereas citrate and isocitrate were reduced (**Fig. 2c**), consistent with altered aconitase activity predicted by the network. Increased pyruvate uptake without changes in lactate secretion, along with reduced TCA intermediate secretion, further suggested rerouting of pyruvate into the TCA cycle (**Fig. 2d**).

**Figure 2.**
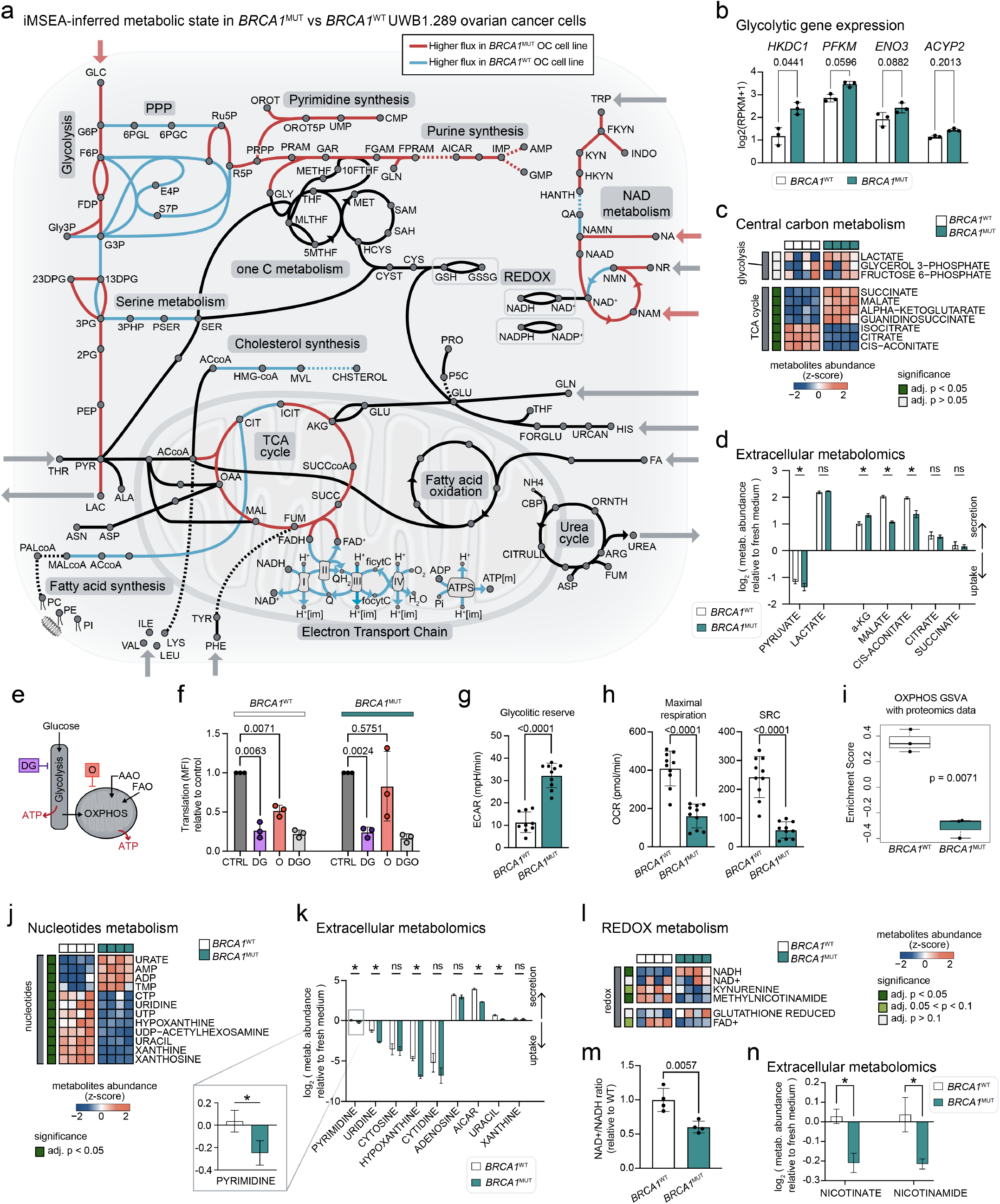
Metabolic state induced by BRCA1-loss in HGSOC UWB1.289 cells. **a**, Representation of metabolic network colored by reaction rates inferred with iMSEA as higher (red) or lower (blue) in *BRCA1*-deficient cells. **b**, Gene expression of glycolytic genes used for the model inference (n=3). **c-d**, Representation of **c**, intracellular and **d**, extracellular metabolites from glycolysis and citric acid (TCA) cycle in *BRCA1*^WT^ and *BRCA1*^MUT^ cells (n=4). **e**, Diagram of the SCENITH assay showing inhibition of glycolysis with 2-deoxyglucose (DG) and oxidative phosphorylation (OXPHOS) with oligomycin (O) to assess ATP production from glucose, amino acid oxidation (AAO), or fatty acid oxidation (FAO). **f**, Metabolic profile of *BRCA1*^WT^ and *BRCA1*^MUT^ cells after SCENITH analysis. Translation level (anti-puro MFI) relative to control is shown (n=3 independent experiments, each in 3-6 replicates). **g**, Impact of *BRCA1*-loss in glycolytic reserve analyzed by glycolytic stress test (n=10). **h**, Quantification by seahorse of maximal respiration and spare respiratory capacity (SRC) in *BRCA1*^WT^ and *BRCA1*^MUT^ cells (n=10). **i**, Enrichment score of oxidative phosphorylation (OXPHOS) pathway calculated by applying GSVA with proteomics data of *BRCA1*^WT^ and *BRCA1*^MUT^ cells (n=3). **j-k**, Representation of **j**, intracellular and **k**, extracellular metabolites from nucleotide metabolism in *BRCA1*^WT^ and *BRCA1*^MUT^ cells (n=4). **l**, Representation of intracellular metabolites from redox metabolism in *BRCA1*^WT^ and *BRCA1*^MUT^ cells (n=4). **m**, NAD^+^/NADH ratio calculated from (MS) metabolomics data (n=4). **n**, Extracellular metabolomics of NAD+ precursors (n=4). Statistical comparisons were performed using a two-tailed t-test unpaired (**b**,**g**,**h**) or paired (**f**), unpaired t-test FDR adjusted for the corresponding peak areas (**c**,**d**,**j**,**k**,**l**,**m**), or Wilcoxon test (**i**). *p<0.05, and n.s. indicates p≥0.05 (not significant). Error bars represent SEM.

To assess glycolytic function beyond basal conditions, we next performed SCENITH analysis^41^, a flow cytometry assay that measures protein synthesis as a surrogate marker of ATP production after inhibition of glycolysis (2-deoxyglucose, DG) or oxidative phosphorylation (oligomycin, O) (**Fig. 2e**). We observed that both cell lines primarily relied on glycolysis to sustain protein translation (**Fig. 2f** and **S3a**). However, *BRCA1*^MUT^ cells exhibited a trend toward greater glycolytic capacity under mitochondrial inhibition, suggesting enhanced metabolic flexibility (**Fig. 2f** and **S3a**). This was further confirmed by glycolytic stress test, which demonstrated a significantly higher glycolytic reserve in *BRCA1*^MUT^ compared to *BRCA1*^WT^ cells (**Fig. 2g** and **S3b**). These results indicate that BRCA1 deficiency confers an increased ability to upregulate glycolysis in response to energetic stress.

Concomitantly, iMSEA predicted a downregulation of mitochondrial oxidative metabolism in *BRCA1*^MUT^ cells. This was supported experimentally by SCENITH analysis, which revealed a trend toward reduced mitochondrial dependence in *BRCA1*^MUT^ cells (**Fig. 2f** and **S3a**), and Seahorse assays, which confirmed lower maximal respiration and reduced spare respiratory capacity in *BRCA1*^MUT^ cells (**Fig. 2h** and **S3c**). Additionally, available proteomic data from the UWB1.289 cell lines^25^ reinforced these observations, as protein set variation analysis showed significantly lower enrichment of the oxidative phosphorylation (OXPHOS) pathway in *BRCA1*^MUT^ cells (**Fig. 2i**). Together, these data demonstrate that *BRCA1*^MUT^ cells exhibit mitochondrial dysfunction and compensate for impaired oxidative metabolism by enhancing glycolytic capacity, especially under stress conditions.

Beyond energy metabolism, iMSEA also predicted increased flux through nucleotide synthesis and NAD metabolism in *BRCA1*^MUT^ cells. Motivated by this prediction, we examined nucleotide metabolism experimentally and although intracellular nucleotide intermediates were lower (**Fig. 2j**), *BRCA1*^MUT^ cells showed increased uptake and decreased secretion of nucleotide precursors (**Fig. 2k**), suggesting enhanced utilization of these substrates for nucleotide biosynthesis. NAD+/NADH homeostasis was also altered, while NAD+ levels were similar across cell types, NADH was significantly higher in *BRCA1*^MUT^ cells, leading to a reduced NAD+/NADH ratio (**Fig. 2l–m**). This redox imbalance likely reflects elevated NADH production from enhanced glycolysis and altered TCA activity, alongside impaired NADH oxidation due to mitochondrial dysfunction. Supporting this, extracellular metabolomics revealed increased uptake of nicotinate and nicotinamide in *BRCA1*^MUT^ cells (**Fig. 2n**), consistent with higher demand for NAD+ precursors to maintain redox balance. Together, these results highlight a reprogramming of nucleotide and redox metabolism in BRCA1-deficient cells.

Overall, BRCA1-deficient UWB1.289 cells exhibit a metabolic shift characterized by reduced mitochondrial oxidative capacity, altered redox balance, and increased glycolytic flexibility under stress. These findings are consistent with prior studies in breast and ovarian cancer, where *BRCA1* expression promoted mitochondrial respiration and suppressed glycolysis^30^, while BRCA1 depletion impaired mitochondrial ATP production and respiratory function in HGSOC cell lines^32^.

Finally, to test whether these metabolic shifts extend beyond the UWB1.289 model, we analyzed proteomic and genetic data from the Cancer Cell Line Encyclopedia (CCLE), which encompasses a diverse panel of ovarian cancer cell lines^42^. Based on published classification by tumor subtype and BRCA1 status^43^, we identified three HGSOC cell lines with *BRCA1* mutation (*BRCA1*^MUT^) and 13 with wild-type *BRCA1* (*BRCA1*^WT^) (**Fig. S4a**). Proteomic analysis of these HGSOC cell lines (available for n=5 *BRCA1*^WT^ and n=2 *BRCA1*^MUT^) revealed enrichment of glycolysis-associated proteins in *BRCA1*^MUT^ cells, whereas *BRCA1*^WT^ cells showed greater abundance of OXPHOS proteins (**Fig. S4b–c**). Although the Wilcoxon test yielded marginal non-significance, the effect size was large with no overlap between groups, and pathway analysis confirmed the trend (**Fig. S4d**). Applying iMSEA to the CCLE proteomics data revealed consistent downregulation of electron transport chain fluxes alongside increased glycolytic flux in *BRCA1*^MUT^ cells (**Fig. S4e–h**). In addition, both purine and pyrimidine synthesis pathways were predicted to carry higher flux in *BRCA1*^MUT^ lines, echoing the results in UWB1.289 cells. The convergence of predictions and experimental data across independent datasets underscores the predictive strength of iMSEA and its value for uncovering BRCA1-associated metabolic alterations in ovarian cancer.

### Task-driven minimal network analysis reveals BRCA1-dependent metabolic reprogramming

While iMSEA captures the global metabolic state of *BRCA1*^WT^ and *BRCA1*^MUT^ cells, it does not directly assess which reaction modules are indispensable for sustaining cellular phenotype-defining functions. To address this, we applied MiNEA^44^ within the iMSEA framework, enabling a task-driven enrichment analysis of minimal subnetworks that support essential metabolic tasks (**Fig. 3a**). Classical enrichment approaches, which typically rely on predefined pathway maps, are limited by fixed pathway boundaries and sparse gene coverage, often leaving large gaps in interpretation. MiNEA overcomes these limitations by identifying minimal subnetworks (MiNs), the smallest sets of reactions required to accomplish a metabolic task, such as the synthesis of a target metabolite. These subnetworks can span multiple canonical pathways, thereby capturing functional modules that are not apparent in database-defined maps. Here, we further strengthened MiNEA by replacing gene-level fold changes with flux fold changes inferred by iMSEA. Unlike gene-level fold changes, which provide incomplete and threshold-dependent coverage, flux values are available for every reaction, allowing us to evaluate not only the transcriptional support but also the functional activity of each minimal network within the overall metabolic state of the cell (**Fig. 3a**).

**Figure 3.**
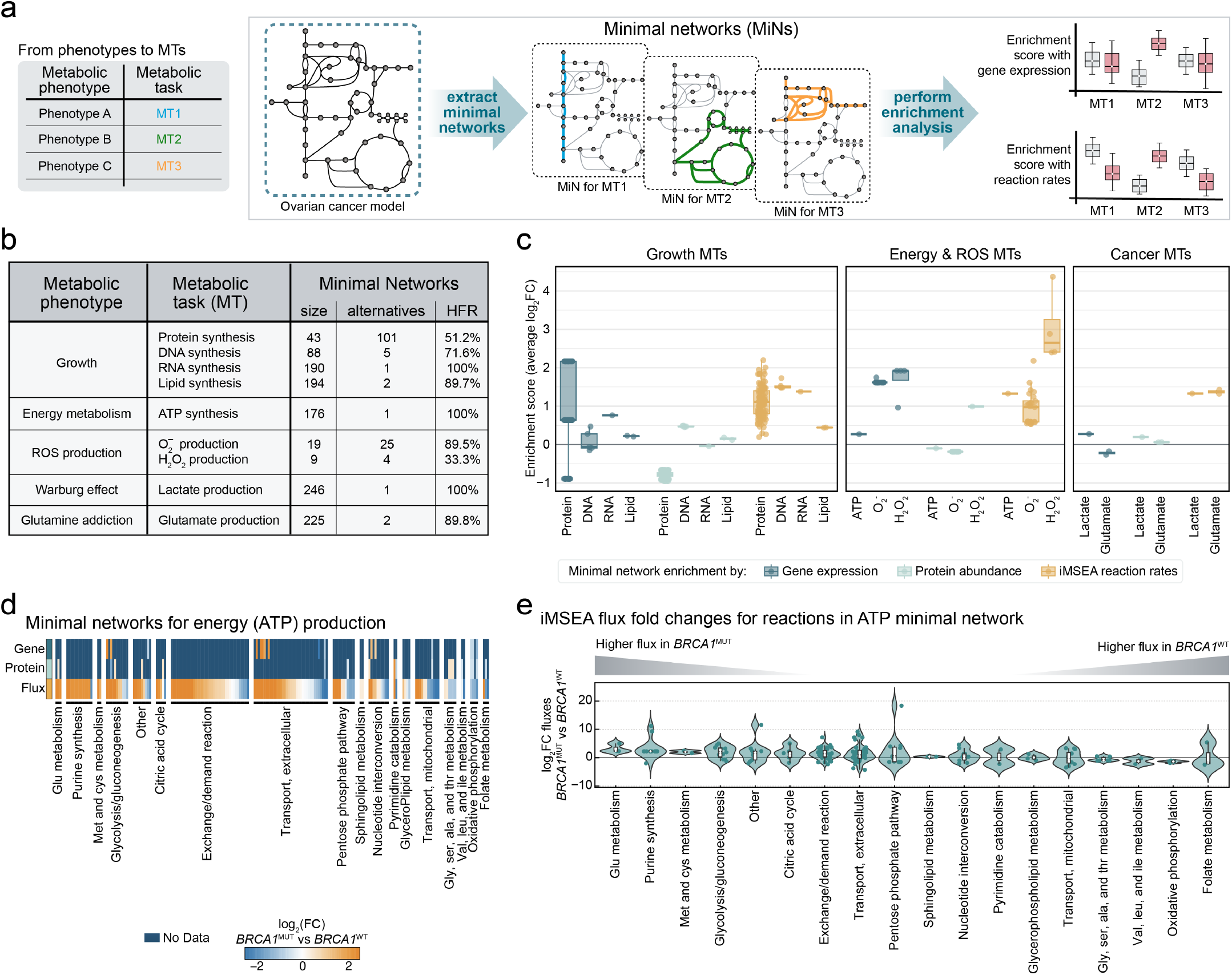
Metabolic minimal networks and enrichment analysis in ovarian cancer cells. **a**, Workflow to represent metabolic phenotypes with the production of metabolic tasks and derive minimal networks (MiNs) from metabolic models and perform enrichment analysis. **b**, Metabolic tasks associated with cancer phenotypes and statistics of the minimal networks: size, number of alternatives, and percentage of common reactions across alternatives. HFR: high-frequency reactions. **c**, Enrichment of metabolic tasks using expression data, proteomics, or the iMSEA in silico fluxes. **d**, Deregulation of reactions required for ATP synthesis when considering gene expression data, proteomics, or inferred iMSEA fluxes. **e**, Subsystem association of reactions present in the minimal network for energy (ATP) production and corresponding fluxes in *BRCA1*^MUT^ vs *BRCA1*^WT^ cells.

Building on this improved enrichment framework, we next applied MiNEA to a curated set of metabolic tasks in *BRCA1*^WT^ and *BRCA1*^MUT^ UWB1.289 cells, chosen to represent three major classes of phenotypic functions (**Fig. 3b**). These included: (i) growth-related tasks, encompassing the synthesis of protein, RNA, DNA, and lipids as essential biomass precursors; (ii) energy and stress-related tasks, defined by the production of ATP, and reactive oxygen species (ROS) such as superoxide anion and hydrogen peroxide; and (iii) cancer-associated tasks, represented by the Warburg effect (lactate secretion) and glutamine addiction (glutamate production). For each task, we used the ovarian cancer metabolic model to identify the minimal set of reactions required to achieve metabolite production, along with alternative subnetworks of equivalent minimal size. These alternatives represent different biochemical routes that can fulfill the same metabolic function. Conservation within each task was then quantified using the high-frequency reaction (HFR) score, which measures the proportion of reactions shared across all alternatives (**Fig. 3b**). Analysis of the MiN properties revealed distinct patterns of metabolic rigidity and flexibility across tasks. Some functions, such as RNA and ATP synthesis or lactate secretion, were supported by large, highly conserved subnetworks with no alternatives, reflecting structural backbones that are consistent across conditions. In contrast, protein synthesis relied on many alternative subnetworks, indicating high pathway plasticity. Other tasks, including lipid and DNA synthesis or glutamate production, displayed an intermediate balance of conservation and flexibility, while ROS-related functions were supported by small, variably conserved networks. These differences suggest that certain metabolic phenotypes are tightly constrained and stable, whereas others are more adaptable, potentially allowing cells to respond to changing demands or stress.

We next investigated how these task-specific subnetworks were differentially utilized in *BRCA1*^MUT^ vs. *BRCA1*^WT^ cells by performing enrichment analysis of the MiNs with three complementary data layers: gene expression, protein abundance, and flux distributions inferred by iMSEA. Across tasks, flux-based enrichment consistently revealed sharper contrasts between the two conditions compared to gene- or protein-based enrichment (**Fig. 3c**). While transcriptional and translational data indicated some deregulation within the MiNs, the average fold changes of significantly altered genes or proteins often remained below threshold, leaving the results largely inconclusive. In contrast, flux-based enrichment revealed a consistent upregulation of activity across all metabolic tasks in *BRCA1*^MUT^ cells, providing a clear signal of global metabolic reprogramming.

To understand the basis for these differences in more detail, we examined the minimal network supporting ATP synthesis (**Fig. 3d**). Gene expression and protein abundance data covered only a subset of the network, limiting interpretability. By contrast, iMSEA-inferred flux values provided complete coverage, enabling a coherent functional assessment of the network. This demonstrates a key advantage of the integrated framework: flux data not only complement but also overcome the sparsity of experimental measurements. Knowledge of the full minimal network further allowed us to explore how specific pathways are differentially used across conditions (**Fig. 3e**). In *BRCA1*^MUT^ cells, flux-based enrichment pointed to elevated activity through glycolysis, the TCA cycle, and purine biosynthesis, consistent with increased anabolic and bioenergetic demands. Conversely, *BRCA1*^WT^ cells showed higher reliance on oxidative phosphorylation and branched-chain amino acid metabolism, suggesting a more respiration-coupled mode of ATP generation. Notably, these pathway-level differences aligned with our experimental validation by SCENITH and Seahorse assays. These results highlight how integrating MiNEA with iMSEA not only improves coverage of minimal networks but also refines our understanding of how distinct pathways are differentially engaged to sustain metabolic functions.

### iMSEA captures metabolic differences in patient tumor data across anatomical sites

The above results demonstrate that iMSEA can accurately capture metabolic states and their architectural features, at least *in vitro*. Through our analysis, we inferred, and validated experimentally, that BRCA1-deficient cell lines downregulate oxidative phosphorylation and likely shift towards glycolysis under stress conditions. However, patient tumors often exhibit a more complex metabolic landscape *in vivo*, shaped by interactions between tumor cells, the microenvironment, and systemic factors, making context-specific analysis critical for understanding therapy response and prognosis.

To assess the applicability of iMSEA for resolving context-specific metabolic states in clinical samples, we applied it to single-cell RNA sequencing (scRNA-seq) data from ovarian cancer patients previously published by Vázquez-García et al.^45^. This dataset includes treatment-naïve HGSOC tumors from a cohort of 42 newly diagnosed patients, with tumor cell transcriptomes annotated across six anatomical sites, including, adnexa, omentum, bowel, peritoneum, upper quadrant, and ascites^45^. We focused on two subgroups as proxies for *BRCA1* status: tumor cells from patients classified as homologous recombination deficient with *BRCA1*-associated tandem duplications (HRD-Dup, n=15; representing *BRCA1*^MUT^) and patients with *CCNE1* amplification-associated foldback inversion bearing tumors (FBI, n=14; representing *BRCA1*^WT^) (**Fig. 4a** and **S5a**). For each anatomical site, we computed gene expression deregulation in cancer cells, and applied iMSEA to infer metabolic fluxes using the ovarian cancer metabolic network (**Fig. 4b** and **S5b**).

**Figure 4.**
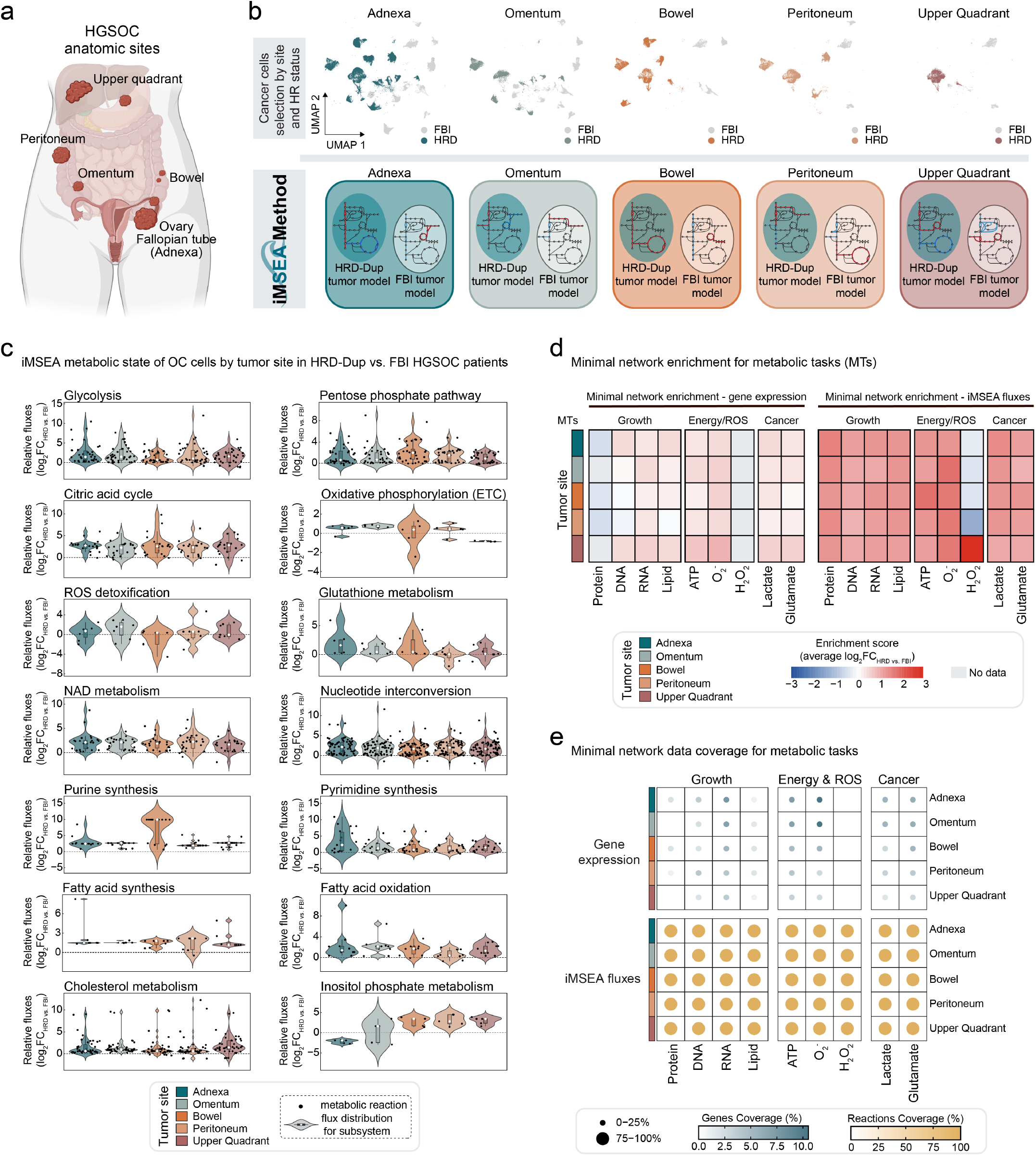
Metabolic state in HGSOC HRD-Dup vs. FBI patients across tumor sites. **a**, Schematic map of anatomical tumor sites. **b**, Workflow scheme: Clustering of cells from different tumor sites (top), and site specific iMSEA inference of metabolic models and states for HRD-Dup and FBI tumors (bottom). **c**, Relative metabolic flux distributions inferred with iMSEA, by tumor site in HRD-Dup vs. FBI HGSOC patients. Log_2_ fold change displayed by reaction and clustered by metabolic subsystem. **d**, Site specific enrichment analysis of ovarian cancer minimal networks using gene expression data or iMSEA fluxes in HRD-Dup vs. FBI patients. **e**, Data coverage of the minimal network composition (genes or reactions) for each metabolic tasks and sites.

The iMSEA analysis revealed distinct metabolic profiles depending on tumor location. HRD-Dup tumors showed across all sites increased inferred-flux in central carbon metabolism (glycolysis, pentose phosphate pathway, and citric acid cycle), NAD and nucleotide metabolism, fatty acid metabolism, fatty acid synthesis, and cholesterol metabolism (**Fig. 4c**), reflecting elevated reliance on these pathways for energy production and macromolecule biosynthesis. Near the primary site (adnexa and omentum), ROS detoxification and glutathione metabolism were elevated in HRD-Dup tumors compared to FBI tumors, suggesting a stronger oxidative stress response in these areas while differences were less pronounced in more distant regions (**Fig. 4c**). Interestingly, the inferred flux through the electron transport chain (ETC) was higher in HRD-Dup tumors closer to the primary site (adnexa, omentum, bowel), but lower in more distant areas like the upper quadrant, where FBI tumors exhibited higher ETC activity (**Fig. 4c**). Conversely, inositol metabolism progressively increased from primary to distant sites in HRD-Dup tumors (**Fig. 4c**), suggesting a potential role in supporting BRCA1-deficient cancer cell survival during metastasis. These findings demonstrate that iMSEA can identify site-specific differences in metabolic activity between tumor subtypes.

While many core metabolic pathways, including central carbon metabolism, nucleotide metabolism, NAD metabolism and fatty acid oxidation, exhibited consistent directional changes between patient-derived tumors and our earlier analysis in cell lines, a few pathways diverged, particularly oxidative phosphorylation, cholesterol metabolism, and fatty acid synthesis. Notably, tumors from metastatic regions like the upper quadrant exhibited a similar phenotype as the cell lines, with an increased activity of central carbon pathways and a reduced flux through mitochondrial oxidative phosphorylation. Interestingly in other HRD-Dup tumor sites, glycolysis and oxidative phosphorylation were simultaneously upregulated, suggesting a metabolic flexibility not captured in cell lines. This dual engagement of energy-producing pathways may reflect adaptation to variable oxygen conditions and other microenvironmental pressures *in vivo*. BRCA1-deficient tumors promote angiogenesis through upregulation of VEGF^25,46^, resulting in increased blood flow and improved access to nutrients and oxygen. We therefore hypothesize that *BRCA1*^MUT^ tumors leverage this vascular advantage to support both increased glycolysis, to sustain rapid proliferation, and elevated oxidative phosphorylation, to meet energy demands *in vivo*. The absence of this dual activation in *in vitro* models may be explained by the uniform nutrient and oxygen availability under culture conditions, which masks the physiological differences present in the tumor microenvironment *in vivo*. Rather than indicating inconsistency, these findings highlight the capacity of iMSEA to detect context-specific metabolic heterogeneity across *in vitro* and *in vivo* settings.

To connect these pathway shifts to functional outcomes, we next applied minimal network enrichment across anatomical sites, following the same iMSEA workflow as in the cell line studies (**Fig. 4d and S5c**). At the transcriptional level, HRD-Dup tumors showed a subtle tendency toward higher expression of task-associated genes compared to FBI tumors. In contrast, flux-based enrichment consistently revealed stronger activity across nearly all tasks in HRD-Dup tumors, with the exception of hydrogen peroxide production. These results parallel our cell line observations and underscore the global upregulation of metabolic tasks associated with BRCA1 deficiency *in vivo*. Analysis of data coverage within the minimal networks revealed that gene expression data encompassed less than 10% of the reactions in most minimal networks (**Fig. 4e**), limiting the interpretability of transcription-based enrichment. By contrast, flux-based inference achieved full reaction coverage, enabling a coherent and systematic assessment of metabolic functions across tasks. This demonstrates that flux-based approaches not only improve coverage but also enable reliable functional interpretation in settings where transcriptional data alone would be insufficient.

Taken together, this comparative analysis shows that iMSEA can reveal both conserved and context-specific features of BRCA1-deficient metabolism. Several metabolic tasks were consistently upregulated in HRD-Dup tumors and in *BRCA1*^MUT^ cell lines, pointing to candidate functional signatures that may serve as translational biomarkers. At the same time, the patient data uncovered site-dependent heterogeneity not captured *in vitro*, emphasizing that knowledge gained from controlled models can be transferred to clinical contexts, but must also be refined in light of the tumor microenvironment. This balance between reproducibility and context-specificity illustrates the strength of iMSEA in bridging mechanistic insight across experimental systems and patient tumors.

### Metabolic reprogramming enables BRCA1-deficient ovarian cancer cells to adapt across metastatic sites

We next asked whether BRCA1-deficient tumors undergo progressive metabolic adaptations as they spread from the adnexal primary site to distant metastases. To investigate this, we analyzed gene expression data from ovarian cancer cells isolated at the primary site (adnexa) and four metastatic sites (omentum, bowel, peritoneum, and upper quadrant) in HRD-Dup tumors^45^ (**Fig. 5a**). Addressing this question required extending iMSEA beyond the pairwise comparisons used earlier (mutant vs. wild-type) to a multi-site framework, where gene expression data from all five anatomical sites were integrated into a unified model of interconnected metabolic networks. Constraining these networks with pairwise gene deregulation enabled us to resolve systematic shifts in reaction activity across metastatic progression (**Fig. 5b**). Analysis of reactions associated with differentially expressed genes in the iMSEA multi-site networks revealed a gradual increase in the number of reactions with significant expression-associated changes from the primary site to more distant metastatic locations (**Fig. S6a-b**). Notably, certain metabolic subsystems, including oxidative phosphorylation, glycolysis, and nucleotide metabolism, were consistently represented among differentially expressed genes across sites (**Fig. S6c**), suggesting that these pathways may play a central role in enabling tumor cells to adapt to the unique metabolic conditions of each metastatic niche.

**Figure 5.**
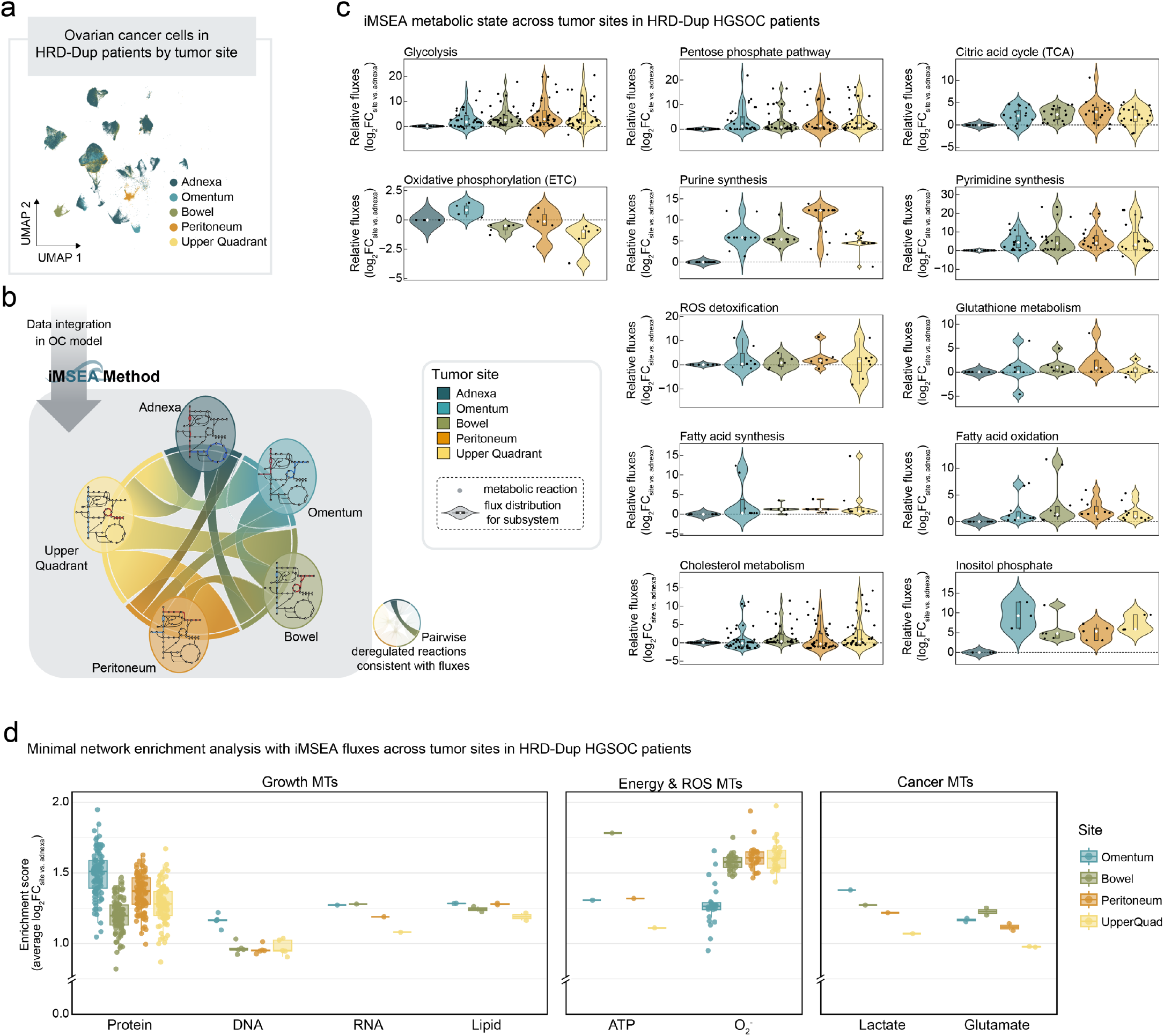
Metabolic heterogeneity across tumor sites in HRD-Dup HGSOC patients. **a**, Ovarian cancer cells from HRD-Dup HGSOC patients segregated by tumor site. **b**, Overview of the data integration simultaneously for five tumor sites (adnexa, omentum, bowel, peritoneum, and upper quadrant). The iMSEA method is used to assess metabolic flux alterations across tumor sites, integrating reaction-level data for metabolic pathway deregulation. **c**, Metabolic pathway activity (log2-transformed flux fold change relative to adnexa) in tumor cells across different sites. ETC: electron transport chain. **d**, Minimal network enrichment analysis for ovarian cancer metabolic tasks (MTs) across sites compared to adnexa.

iMSEA computational analysis of metabolic flux distributions revealed broadly increased central carbon metabolism in metastatic regions compared to the adnexal primary site. Glycolysis, the pentose phosphate pathway, and the citric acid cycle all carried higher flux, accompanied by increased nucleotide synthesis (purine and pyrimidine pathways), consistent with elevated biosynthetic demand (**Fig. 5c**). ROS detoxification and glutathione metabolism were particularly enhanced in peritoneal and bowel metastases, suggesting higher oxidative stress in these microenvironments. Fatty acid synthesis and oxidation also increased across metastatic sites, whereas cholesterol metabolism remained comparatively stable. Inositol phosphate metabolism displayed site-dependent variability, and oxidative phosphorylation fluxes showed marked divergence between sites, highlighting microenvironment-specific pressures on mitochondrial function (**Fig. 5c**).

To translate these pathway-level findings into core metabolic functions to sustain tumor growth and survival, we performed flux enrichment of minimal networks across metastatic sites relative to the adnexal primary tumors (**Fig. 5d** and **S6d**). This revealed upregulation of growth-related tasks, particularly protein synthesis, across all metastatic locations, highlighting the enhanced biosynthetic demands of disseminated BRCA1-deficient cells. Energy- and ROS-associated tasks also increased in a site-dependent manner, supporting bioenergetic needs and adaptation to local metabolic stress. Cancer hallmark functions, such as lactate secretion and glutamate production, were consistently enhanced, reinforcing glycolytic and glutamine-dependent phenotypes.

Consistent with previous reports of elevated central carbon metabolism, oxidative stress adaptation, and lipid transfer–driven β-oxidation in ovarian cancer metastases^47-49^, our analysis indicates that BRCA1-deficient HGSOC tumors disseminating from the adnexa undergo metabolic reprogramming, producing tumor populations that are highly metabolically active, stress-adapted, and potentially more aggressive.

## DISCUSSION

Metabolic reprogramming is a hallmark of cancer and other metabolic diseases, and a major determinant of progression, therapy response, and disease heterogeneity^1,50,51^. Yet, despite the wealth of transcriptomic and proteomic data available, conventional enrichment approaches that operate directly on omics profiles are limited in their ability to resolve how these changes alter functional metabolism. They typically fail to capture flux directionality, stoichiometric constraints, or the emergence of bottlenecks that define metabolic vulnerabilities. Addressing this gap requires methods that can systematically link gene expression shifts to the biochemical logic of cellular networks.

To meet this need, we developed iMSEA (*in silico* Metabolic State and Enrichment Analysis), a systems biology framework that embeds omics changes into genome-scale metabolic networks and infers flux-level activity. iMSEA integrates transcriptomic data into constraint-based models and uses flux balance analysis combined with mixed-integer linear programming to estimate directional flux changes across reactions. By performing enrichment on the fold changes of these inferred fluxes, iMSEA directly connects expression shifts to their functional metabolic consequences, while accounting for reaction stoichiometry, cofactor availability, and subsystem context. This provides a level of mechanistic resolution not achievable by conventional enrichment, enabling the identification of context-dependent flux alterations and potential vulnerabilities. Importantly, iMSEA is not limited to a specific disease or dataset: it can incorporate bulk RNA-seq, proteomics, or single-cell RNA-seq, and can be applied to reconstructions ranging from curated networks^13,15,16^ to full genome-scale models^5,6^.

As a proof of principle, we applied iMSEA to BRCA1-deficient high-grade serous ovarian cancer (HGSOC), where metabolic rewiring contributes to tumor progression and therapy resistance. In vitro, BRCA1-deficient HGSOC cell lines displayed enhanced flux through glycolysis, nucleotide synthesis, and the TCA cycle, coupled with reduced oxidative phosphorylation, fatty acid synthesis, and cholesterol metabolism. In contrast, patient-derived single-cell RNA-seq revealed a broader metabolic upregulation, including glycolysis, oxidative phosphorylation, and lipid metabolism. This discordance reflects well-established differences between cell culture and tumors *in vivo*, including stromal contributions, nutrient gradients, and inter-patient heterogeneity. Rather than a limitation, this highlights iMSEA’s sensitivity to context-specific metabolic states and underscores the importance of model selection when translating results between experimental systems and patients.

Beyond pairwise comparisons, iMSEA is well suited for analyzing multiple metabolic states simultaneously. In metastatic tumors, for example, we used iMSEA to resolve progressive adaptations across anatomical sites, revealing how local environments drive site-specific metabolic programs. This capacity to dissect metabolic heterogeneity across space, time, or treatment conditions extends iMSEA’s utility beyond static snapshots, providing a dynamic view of metabolic evolution.

By resolving flux-level activity across genotypes, tumor types, and microenvironments, iMSEA establishes a foundation for studying how metabolic plasticity shapes tumor behavior, metastatic fitness, and therapy response^52-54^. Its generality makes it broadly applicable to oncology and metabolic medicine, where context-specific rewiring is a recurring theme. Integration with spatial metabolomics, isotope tracing, or functional perturbations will further enhance its capacity to reveal how metabolic programs confer survival advantages, modulate stromal crosstalk, or enable immune evasion^55^.

In summary, iMSEA provides a generalizable platform for translating omics data into mechanistically grounded metabolic insight. By bridging gene expression changes with flux-level consequences, it enables the discovery of context-specific adaptations and vulnerabilities. While our BRCA1-deficient HGSOC analysis illustrates its ability to capture biologically meaningful rewiring *in vitro, in vivo*, and across metastatic niches, the framework extends well beyond this case. iMSEA thus offers a versatile strategy to interrogate metabolic heterogeneity in cancer and other diseases, supporting efforts to target the adaptive programs that sustain pathologic metabolism.

## DATA AND CODE AVAILABILITY

All input files, processed data, and scripts used for running iMSEA, including differential expression thresholds, optimization settings, and final reaction sets, are available on GitHub (https://github.com/mariamasid/iMSEA).

## AUTHOR CONTRIBUTIONS

Conceptualization, M.M., V.H., G.C., D.D.L.; Methodology, Software, and Formal Analysis, M.M.; Investigation, M.M., I.A.R, D.B, F.D.C., M.D., P.G., Y.O.M.; Resources, V.H., G.C., D.D.L., N.V., D.Z., S.P.S.; Visualization, M.M.; Writing – Original Draft, M.M.; Writing – Review & Editing, M.M., V.H., G.C., D.D.L. wrote the manuscript and all authors reviewed and approved the final version; Project Administration, M.M.; Supervision, V.H., G.C., D.D.L.; Funding Acquisition, V.H., G.C., D.D.L.

## ACKNOWLEDGMENTS

We thank Dr. Matteo Morotti and Dr. Eleonora Ghisoni for constructive discussions; and the Metabolomics Platform team at the Faculty of Biology and Medicine, University of Lausanne for metabolome analysis. This work was generously supported by the Ludwig Institute for Cancer Research to G.C., by a Swiss National Science Foundation (SNF) advanced grant to G.C. (216381), by the Cancera and Mats Paulson Foundations to G.C., by the Ludwig Institute for Cancer Research (Myeloid Cells in Cancer Initiative, MCCI project) to D.D.L., by the DOD OCA Early Career Investigator (ECI) W81XWH2210703 Award OC210038 to D.D.L, by the Swiss National Science Foundation (SNSF) Sinergia project CRSII5_198543 to V.H., and by the École Polytechnique Fédérale de Lausanne (EPFL) to V.H.

## MATERIALS AND METHODS

### CELL CULTURE

UWB1.289 (BRCA1^MUT^) and UWB1.289+BRCA1 (BRCA1^WT^) cell lines were obtained from ATCC and cultured as indicated by manufacturer. The UWB1.289 cell line was cultured in a 50% ATCC-formulated RPMI-1640 Medium (#30-2001) and 50% MEGM Bullet kit (Mammary Epithelial Growth Medium from Clonetics/Lonza #CC-3150) supplemented with 3% heated-inactivated fetal bovine serum, 100µg/mL penicillin and 100µg/mL streptomycin at 37^°^C in 5% CO_2_. The UWB1.289+BRCA1 cell line was cultured in the same media plus 200µg/mL G-418.

### CANCER CELL LINE ENCYCLOPEDIA DATA

Proteomics, mutation, and copy number variation data from the Cancer Cell Line Encyclopedia (CCLE) were utilized to investigate metabolic pathway deregulations in BRCA1-deficient high-grade serous ovarian cancer (HGSOC) cell lines. Sixteen HGSOC cell lines were selected based on subtype classification^43^. Differential protein abundance and pathway enrichment analyses were performed using the limma^56^ and GSVA^57^ R packages, respectively. Statistical significance was assessed with Wilcoxon rank-sum tests.

### COHORT OF HGSOC PATIENTS

We analyzed publicly available single-cell RNA sequencing (scRNA-seq) data from a cohort of 42 newly diagnosed, treatment-naïve high-grade serous ovarian cancer patients originally reported by Vazquez-Garcia et al^45^. Patients were classified into homologous recombination deficient (HRD-Dup, n=16) and homologous recombination proficient (FBI, n=14) groups. Pre-processed Seurat objects with cells annotated into nine major lineages were retrieved (available via synapse with accession number syn25569736) and used for downstream analyses. Pathway and hallmarks enrichment analysis for patient pseudo-bulk data were inferred with the GSVA^57^ R package and statistical analyses were done using Wilcoxon rank-sum tests. Differential gene expression for patient pseudo-bulk data was calculated with the limma R package^56^, and the Wilcoxon rank-sum test was used for statistical significance.

### WESTERN BLOT

BRCA1 protein expression in UWB1.289 and UWB1.289+BRCA1 cell lines was analyzed using SDS-PAGE and Western blot. Nuclear protein extracts were obtained from cells growing in logarithmic phase using NE-PER Nuclear and Cytoplasmic Extraction Reagents (Thermo Fisher, 78835). Protein concentration was determined using the Quick Start™ Bradford 1× Dye Reagent (Bio-Rad, 5000205). Nuclear protein extracts (50ug) were loaded into a NuPAGE™ 3–8% Tris-Acetate Gel (Invitrogen, EA0375PK2) and resolved by gel electrophoresis in Tris-Acetate SDS Running Buffer (Novex, LA0041). PageRuler Plus Prestained Protein Ladder (Thermo Fisher, 26619) was used as a molecular weight marker.

Proteins were transferred to a 0.2 µm nitrocellulose membrane (Bio-Rad, 1620112) using NuPAGE™ Transfer Buffer (Novex, NP0006-1). The membrane was blocked in 5% non-fat dry milk (Millipore Sigma, 70166) in Tris-buffered saline (TBS)-Tween 0.05% (TBST), followed by overnight incubation at 4°C with a mouse antibody anti-hBRCA1 (MS110; Millipore Merck, MABC199, 1:1000) in 5% milk/TBST. The primary antibody was detected using a secondary horseradish peroxidase (HRP)-conjugated goat anti-mouse antibody (DAKO, P0447). β-ACTIN was used as loading control (Santa Cruz, sc-47778). Nitrocellulose membrane was developed using Azure Radiance Plus (Fentogram HRP substrate, AC2103) and visualized using the Fusion FX Imaging System (Vilber). Densitometric analysis was performed using ImageJ software.

### CELL COUNT

UWB1.289 and UWB1.289+BRCA1 cell lines were plated in 6 well plates at 150,000 cells per well in triplicate. After 47 hours from cell adhesion, cells growing in logarithmic phase were detached using Trypsin and counted by trypan blue dead cells exclusion using LUNA-FX7™ Automated Cell Counter (LOGOS BIOSYSTEMS). Cell doubling time was calculated using the following equations:

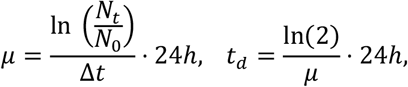

where, *μ* is the growth rate, Δ*t* are the hours of growth, *N*_0_ are the number of cells seeded, *N*_*t*_ are the number of cells harvested, and *t*_*d*_ represents the doubling time. The experiment was repeated twice.

### GROWTH ASSAY

UWB1.289 and UWB1.289+BRCA1 were prepared at a dilution of 50,000 cells/ml and seeded at 100ul/well in triplicate. For day 0 detection, 10ul per well of CCK-8 solution was added 4 hours after cells seeding, and the absorbance at 450nm was read after 2 hours using a microplate reader (SPARK Multimode Microplate Reader, Tecan). Cell growth was monitored every day until day 4. Culture media was changed after 48h. The experiment was repeated three-times.

### UWB1.289 CELL LINES RNA SEQUENCING DATA ANALYSIS

Transcriptomics profiling by bulk RNA sequencing data from UWB1.289 (n=3) and UWB1.289+BRCA1 (n=3) cell lines were collected from the work by Bruand *et al*. (GEO: GSE120792; see original manuscript^25^ for information on data processing). Gene set variation analysis was assessed with the GSVA R package^57^ and differential gene expression analysis was performed using the regularized linear model as implemented in the limma R package^56^.

### UWB1.289 CELL LINES PROTEOMICS ANALYSIS

LC-MS/MS proteomics data of the cell lines were obtained from the work by Bruand et al^25^. Protein expression was also subjected to variation analysis and differential abundance analysis in the same way as described for transcriptomics analyses.

### RECONSTRUCTION OF AN OVARIAN CANCER METABOLIC MODEL

A metabolic model for the study ovarian cancer was reconstructed using the redHUMAN method^17^ applied to the thermodynamically curated human genome-scale metabolic model Recon3D^5,17^. The redHUMAN approach extracts relevant metabolic subsystems from the genome-scale network while preserving key reactions involved in nutrient uptake, biomass synthesis, and metabolic interactions. We selected 11 core subsystems: glycolysis/gluconeogenesis, citric acid cycle, pentose phosphate pathway, glutamate metabolism, reactive oxygen species detoxification, glycine/serine/alanine/threonine metabolism, urea cycle, arginine and proline metabolism, purine synthesis, pyrimidine synthesis, and oxidative phosphorylation. Nutrient availability was defined based on RPMI medium composition in the extracellular compartment. Model reduction parameters used included D=1 for redGEM, Smin for redGEMX, and Sminp3 for lumpGEM. Lumped reactions generated by lumpGEM were explicitly incorporated into the model. The resulting model comprises 1,099 genes, 2,369 reactions, and 974 metabolites.

### iMSEA WORKFLOW – METABOLIC STATE

The iMSEA (in-silico metabolic state and enrichment analysis) computational workflow was designed to infer metabolic states associated with specific gene deregulation profiles. It integrates gene expression data into a metabolic network, constraining reaction fluxes based on observed transcriptional changes. By systematically adjusting flux distributions, iMSEA enables the identification of metabolic pathways that are functionally impacted under different conditions. It encompasses the following steps:

#### 1. Defining physiological conditions and metabolic model contextualization

The user can provide physiological information, such as growth rates, extracellular medium composition, nutrient uptake/secretion rates, or known reaction fluxes, to refine the metabolic model. While these constraints are not all required, incorporating them can enhance the accuracy of the metabolic state predictions.

#### 2. Mapping Genes to Reactions

Gene expression data is mapped to metabolic reactions using gene-protein-reaction (GPR) rules, which define the relationships between genes, enzymes, and reactions in the network. The user needs to specify a threshold for the fold-change (FC) of genes (e.g., a threshold of 1.3 will classify genes as upregulated if FC>1.3, and downregulated if FC<1/1.3) and one or more significance thresholds (e.g., p-values of 0.01, 0.05, 0.1) to identify differentially expressed genes (DEGs) at different confidence levels.

For reactions regulated by multiple genes, iMSEA applies standard systems biology rules^9-11^:

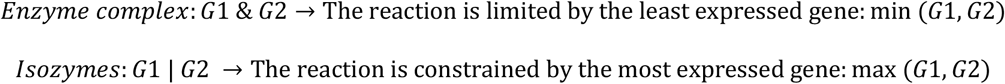

iMSEA then determines the corresponding differentially expressed reactions for each selected threshold, allowing for a flexible and graded integration of gene expression changes into the metabolic network.

#### 3. Integrating Gene Expression into the Metabolic Network

To incorporate the gene expression changes into the metabolic model, iMSEA applies the REMI (Relative Expression and Metabolomic Integrations) method^14^, which constrains reaction flux ratios based on gene expression fold changes. In iMSEA this process is performed sequentially, prioritizing reactions associated with genes that have the most significant deregulation.

Therefore, for a pair of conditions, *a, b*, having identified the set of deregulated reactions for the specific fold change and p value thresholds, we define the set of upregulated reactions 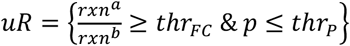 and the set of downregulated reactions 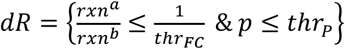.

Then, the following mixed integer linear programming (MILP) problem is formulated to maximize the number of reaction rates with flux ratios between both conditions consistent with the deregulation profile, while satisfying stoichiometric and thermodynamic constraints:

*objective function* 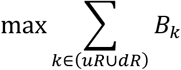

*TFA constraints*

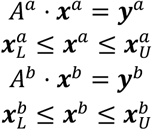

*flux ratio constraints*

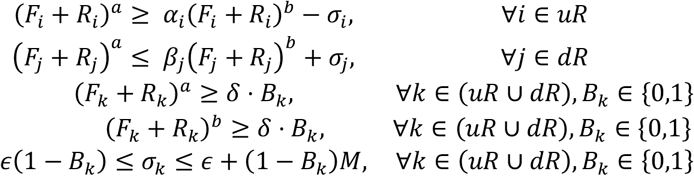

where ***A***^***a***^, ***x***^***a***^, ***y***^***a***^ are the thermodynamics-based flux analysis (TFA) matrix, vector of variables and right-hand side term for condition *a*, and ***A***^***b***^, ***x***^***b***^, ***y***^***b***^ are the corresponding for condition *b*. The TFA formulation includes quasy-steady state constraints (***S* ⋅ *v*** = **0**) and thermodynamic constraints (Δ***G***’), with reaction fluxes, metabolite concentrations, and thermodynamic variables being optimized simultaneously (see ^58^ for full formulation details). In the TFA formulation, reaction fluxes are split into the net-forward and net-reverse fluxes, so that, *v*_*i*_ = *F*_*i*_ ™ *R*_*i*_, where *F*_*i*_, *R*_*i*_ ≥ 0 ensuring only one is active at a time. In the flux-ratio constraints, *α*_*i*_, *β*_*i*_ represet the gene expression fold change ratios for reaction *i. δ* is a user-defined basal flux, while *B*_*k*_ is a binary variable that determines whether reaction k follows the expected expression pattern. In this case if *B*_*l*_ = 1, then the flux through reaction k satisfies the expression fold change. The slack variable *σ*_*i*_ allows for minor inconsistencies, bounded by a small constant *ϵ* and a large constant *M*. In this study, *δ* = 1.5 ⋅ 10^-4^, *ϵ* = 10^-5^, *M* = 1000 and CPLEX12.10 solver was used for solving the MILP problem.

Although the solver guarantees an optimal solution, it may not be unique, as alternative solutions may exist. To account for this, iMSEA iteratively identifies alternative optimal solutions by enforcing the following constraints after obtaining the first solution:

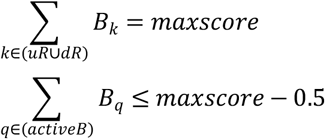

here *maxscore* is the total number of reactions satisfying the expression constraints in the first solution, and *activeB* represents the set of consistent reactions in that solution, i.e., *B*_*q*_ = 1 ∀*q* ∈ *activeB*.

The reactions consistently active across alternative solutions are retained, and the process is repeated for the next p-value threshold. To ensure that the most significant deregulated reactions influence metabolic flux predictions the most, constraints derived from lower p-value thresholds are kept in subsequent iterations, while additional reactions are incorporated at higher p-value thresholds.

While this formulation describes two conditions (*a, b*) and a single ratio (*a*/*b*), it is readily extended to multiple conditions (*a, b, c*, …) and the corresponding pairwise ratios (*a*/*b, a*/*c, b*/*c*, …). The user selects which ratios to integrate based on their experimental design.

#### 4. Flux Variability Analysis

Flux variability analysis^59^ is applied to determine the range of possible flux values for each reaction while satisfying all model constraints, including stoichiometric, thermodynamic, and gene-expression constraints. In this step, the reactions identified as consistent in the above iMSEA optimization are fixed, ensuring that their flux ratios remain constrained according to gene expression-derived limits. For all reactions in the network, two optimization problems are solved to determine its minimum and maximum allowable flux. This analysis provides insight into the flexibility of metabolic fluxes under the imposed constraints.

#### 5. Flux Sampling and Distribution Analysis

Due to the constraints imposed on flux ratios by gene expression data, the space of feasible solutions in iMSEA becomes non-convex, as some bidirectional reactions (BDRs) can carry basal flux in either the forward or reverse direction. Since the Artificial Centering Hit-and-Run (ACHR) Markov Chain Monte Carlo (MCMC) algorithm, implemented in the COBRA toolbox^60^, is designed for convex spaces, iMSEA employs a partitioning strategy to enable efficient flux sampling.

First, the set of bidirectional reactions constrained by gene-expression data is identified. Then, to generate representative samples across the entire feasible solution space, iMSEA iteratively partitions the space into convex subspaces by selectively fixing the directionality of an arbitrary subset of these BDRs. In each iteration, one BDR is randomly selected and temporarily fixed to a forward or reverse direction. A feasible solution is computed under this partial directionality assignment, and the remaining BDRs are fixed according to the directions realized in this solution. This process defines a unique convex subspace. Within each of these subspaces, ACHR-MCMC sampling is performed while preserving all stoichiometric, thermodynamic, and REMI-derived constraints. This process is repeated across multiple subspaces, ensuring that both forward and reverse states of each BDR are sampled.

By systematically covering all convex subspaces, iMSEA reconstructs a representative distribution of metabolic flux states while accurately reflecting the constraints derived from gene expression data. This approach allows for a comprehensive assessment of reaction activity under different conditions, capturing both tightly constrained and more flexible metabolic flux patterns.

### iMSEA WORKFLOW – MINIMAL NETWORK ENRICHMENT ANALYSIS

#### 1. Formulation of Metabolic Tasks for Cancer Phenotypes

Using the MiNEA framework^44^, five metabolic phenotypes relevant to cancer cells were represented as metabolic tasks. These included proliferation, energy metabolism, ROS production, Warburg effect, and glutamine addiction. Each task was defined by the production of specific metabolites: (1) Proliferation: production of macromolecules via the biomass reaction; (2) Energy metabolism: ATP production; (3) Oxidative stress: superoxide anion and hydrogen peroxide production; (4) Warburg effect: lactate production; and (5) Glutamine addiction: glutamate production

#### 2. Generation of Minimal Networks (MiNs)

For each metabolic task, we used the ovarian cancer model to generate minimal networks (MiNs), defined as the smallest set of reactions required to achieve the task. Each MiN included intracellular reactions, transport reactions, and boundary reactions necessary for the production of task-specific metabolites.

#### 3. Minimal Network Enrichment Analysis

Deregulation within each MiN was assessed using the MiNEA (Minimal Network and Enrichment Analysis) framework^44^. Gene-protein-reaction (GPR) rules were applied to integrate multi-omics data, including gene expression, proteomics, and flux distributions inferred via iMSEA. A deregulation score was assigned to each MiN based on the combined activity of its constituent reactions, enabling the identification of differentially regulated metabolic subnetworks across *BRCA1*^WT^ and *BRCA1*MUT conditions.

### APPLICATION OF THE iMSEA WORKFLOW

#### In vitro UWB1.289 cell lines

The iMSEA workflow was applied to compare *BRCA1*-mutant (*BRCA1*^MUT^) and *BRCA1*-wildtype (*BRCA1*^WT^) UWB1.289 cell lines (n=3 per group). Differentially expressed genes (DEGs) were identified using a fold-change threshold of 1.3 and four significance levels (p < 0.05, 0.1, 0.3, and 0.5). These DEGs were mapped to metabolic reactions, and reaction fluxes were constrained accordingly. A total of 725 reactions were found consistent with the deregulation profile.

#### CCLE cell lines

The iMSEA workflow was applied to *BRCA1*^MUT^ (n=3) and *BRCA1*^WT^ (n=13) ovarian cancer cell lines from the CCLE. Proteomics data were used to identify differentially abundant proteins (DAPs) with a fold-change threshold of 1.3 and four significance levels (p < 0.05, 0.1, 0.3, 0.5). DAPs were mapped to metabolic reactions, and iMSEA constrained reaction fluxes accordingly, identifying 465 consistent reactions.

#### Patient data from primary and metastatic ovarian cancer

The iMSEA workflow was applied to compare *BRCA1*^MUT^-like (HRD-Dup, n=16) and *BRCA1*^WT^-like (FBI, n=14) tumors across primary adnexal lesions and four metastatic sites (omentum, peritoneum, bowel, upper quadrant). Gene expression data were pseudo-bulked per patient, and DEGs were identified with a 1.2-fold-change threshold and up to four significance levels (p < 0.05, 0.1, 0.3, 0.5). Biologically relevant doubling times of 75 and 90 days^61^ were used to constrain growth rates. A total of 46 consistent reactions were identified in adnexa, 36 in omentum, 55 in bowel, 57 in peritoneum, and 77 in upper quadrant samples.

#### Comparative metabolic states across sites

iMSEA was used to compare metabolic flux profiles of *BRCA1*^MUT^-like tumors across five anatomical sites: adnexa (n=13), omentum (n=9), bowel (n=6), peritoneum (n=4), and upper quadrant (n=2). Data were pseudo-bulked per patient, and DEGs were identified with a 1.2-fold-change threshold and three significance levels (p < 0.05, 0.1, 0.3). Gene expression was mapped to reactions, and flux distributions adjusted to identify shared and unique metabolic states. Biologically relevant doubling times of 75 and 90 days^61^ were used to constrain growth rates. A total of 278 consistent reactions were identified across the sites.

### METABOLOMICS

Cells from UWB1.289 and UWB1.289+BRCA1 lines were seeded at 0.5 × 10^6^ cells per 60 mm dish (n=4) and cultured for 48 hours. Spent media and cell pellets were collected after 48h and stored at ™80°C. Fresh media samples were stored at -80°C on the day of cell seeding.

Cell lysates were extracted with 80% methanol, homogenized, centrifuged, evaporated to dryness, and reconstituted in methanol:water (4:1) according to total protein content. Media samples were extracted with methanol and centrifuged. Extracts were analyzed by ultra-high performance liquid chromatography tandem mass spectrometry (UHPLC-MS/MS) in multiple reaction monitoring mode, using a Triple Quadrupole mass spectrometer. Both positive and negative electrospray ionization modes were employed to maximize metabolome coverage^62,63^. Data acquisition was performed with appropriate software, and quality control (QC) samples were run periodically to monitor stability and correct signal drift. Metabolites exceeding 20% coefficient of variation in QC samples were excluded from analysis^64^.

### GLYCOLYTIC STRESS TEST

UWB1.289 and UWB1.289+BRCA1 cells were plated in a Seahorse microplate (5x10^4^ cells/well) with corresponding culture media for 18 hours before the experiment. The day of the experiment cells were preincubated at 37^0^C for 45 min in the absence of CO_2_ in Seahorse media supplemented with pyruvate (1mM) and glutamine (2mM). ECAR was measured under basal conditions and after the addition of the following compound and drugs: glucose (10mM), oligomycin (1mM), and 2-deoxy-D-glucose 50mM, following standard glycolytic stress test protocol.

### SEAHORSE

UWB1.289 and UWB1.289+BRCA1cell oxygen consumption rate (OCR) was measured using a Seahorse XF96 Extracellular Flux Analyzer. Cells were plated in a Seahorse microplate (5x10^4^ cells/well) with corresponding culture media for 18 hours before the experiment. The day of the experiment cells were preincubated at 37^0^C for 45 min in the absence of CO_2_ in Seahorse media supplemented with glucose (10mM), pyruvate (1mM) and glutamine (2mM). OCR was measured under basal conditions and after the addition of the following drugs: oligomycin (1mM), carbonyl cyanide-p-trifluoromethoxyphenylhradone (FCCP, 1.5mM) and rotenone/Antimycin A (0.5mM), following standard mitostress protocol.

### SCENITH ANALYSIS

Cells from the UWB1.289 and UWB1.289+BRCA1 cell lines were cultured as specified by manufacturer. On the day of the assay, cells were trypsinized, washed, counted and resuspended in RPMI media without methionine at a concentration of 150.000 in 200ul per well of a 48well plate. Cells were incubated at 37^°^C for 30min to 1hour. Four or five experimental replicates per cell line and per treatment were prepared. Cells were treated for 60min at 37^°^C with 2-Deoxy-D-Glucose (DG, final concentration 100mM), Oligomycin (Oligo, final concentration 1 μM), or a combination of the two drugs. Untreated cells were also included as a control group for assessing the total protein synthesis. Puromycin (at a final concentration of 10μg/mL) was added during the last 30min of the metabolic inhibitors treatment. As a negative control, untreated cells with no puromycin were used. After puromycin treatment, cells were washed with ice-cold PBS, Fc receptors were blocked using an Fc receptor binding inhibitor and stained with Aqua fluorescent cell viability marker for 30min in PBS. After washing, cells were fixed and permeabilized using the FOXP3 fixation kit. Intracellular staining of puromycin using an anti-Puro Alexa Fluor 647 antibody was performed by incubating cells for 1h at 4^°^C. Fluorescently labeled cells were analyzed immediately after staining by a Fortessa flow cytometry analyzer (BD Biosciences). The median florescence intensity of puromycin was measured within the live cells population across all conditions. SCENITH calculations for glucose and mitochondrial dependence, as well as fatty acid and amino acid oxidation and glycolytic capacity were performed as specified in the original study^41^.

### STATISTICAL ANALYSES AND REPRODUCIBILITY

All statistical analyses were performed using R (v4.4.1) and GraphPad Prism (v10.1.1). Metabolomics data were analyzed using MetaboAnalyst (v6.0). All statistical details are provided in the figures, and Results section. Experiments were conducted with both biological and technical replicates, as specified in the figure legends, to ensure reproducibility. The number of independent replicates (n) is reported for each experiment.

## REFERENCES

1. Hanahan, D. & Weinberg, R. A. Hallmarks of Cancer: The Next Generation. Cell 144, 646–674 (2011). 10.1016/j.cell.2011.02.013

2. Hanahan, D. Hallmarks of Cancer: New Dimensions. Cancer Discov 12, 31–46 (2022). 10.1158/2159-8290.Cd-21-1059

3. Thiele, I. & Palsson, B. O. A protocol for generating a high-quality genome-scale metabolic reconstruction. Nat Protoc 5, 93–121 (2010). 10.1038/nprot.2009.203

4. Gu, C. D., Kim, G. B., Kim, W. J., Kim, H. U. & Lee, S. Y. Current status and applications of genome-scale metabolic models. Genome Biol 20 (2019). 10.1186/s13059-019-1730-3

5. Brunk, E. et al. Recon3D enables a three-dimensional view of gene variation in human metabolism. Nat Biotechnol 36, 272–281 (2018). 10.1038/nbt.4072

6. Robinson, J. L. et al. An atlas of human metabolism. Sci Signal 13 (2020). 10.1126/scisignal.aaz1482

7. O’Brien, E. J., Monk, J. M. & Palsson, B. O. Using Genome-scale Models to Predict Biological Capabilities. Cell 161, 971–987 (2015). 10.1016/j.cell.2015.05.019

8. Yizhak, K., Chaneton, B., Gottlieb, E. & Ruppin, E. Modeling cancer metabolism on a genome scale. Mol Syst Biol 11 (2015). 10.15252/msb.20145307

9. Becker, S. A. & Palsson, B. O. Context-specific metabolic networks are consistent with experiments. Plos Comput Biol 4 (2008). 10.1371/journal.pcbi.1000082

10. Zur, H., Ruppin, E. & Shlomi, T. iMAT: an integrative metabolic analysis tool. Bioinformatics 26, 3140–3142 (2010). 10.1093/bioinformatics/btq602

11. Jerby, L., Shlomi, T. & Ruppin, E. Computational reconstruction of tissue-specific metabolic models: application to human liver metabolism. Mol Syst Biol 6 (2010). 10.1038/msb.2010.56

12. Agren, R. et al. Reconstruction of Genome-Scale Active Metabolic Networks for 69 Human Cell Types and 16 Cancer Types Using INIT. Plos Comput Biol 8 (2012). 10.1371/journal.pcbi.1002518

13. Agren, R. et al. Identification of anticancer drugs for hepatocellular carcinoma through personalized genome-scale metabolic modeling. Mol Syst Biol 10 (2014). 10.1002/msb.145122

14. Pandey, V., Hadadi, N. & Hatzimanikatis, V. Enhanced flux prediction by integrating relative expression and relative metabolite abundance into thermodynamically consistent metabolic models. Plos Comput Biol 15 (2019). 10.1371/journal.pcbi.1007036

15. Wang, Y. L., Eddy, J. A. & Price, N. D. Reconstruction of genome-scale metabolic models for 126 human tissues using mCADRE. Bmc Syst Biol 6 (2012). 10.1186/1752-0509-6-153

16. Vlassis, N., Pacheco, M. P. & Sauter, T. Fast Reconstruction of Compact Context-Specific Metabolic Network Models. Plos Comput Biol 10 (2014). 10.1371/journal.pcbi.1003424

17. Masid, M., Ataman, M. & Hatzimanikatis, V. Analysis of human metabolism by reducing the complexity of the genome-scale models using redHUMAN. Nat Commun 11, 2821 (2020). 10.1038/s41467-020-16549-2

18. Machado, D. & Herrgård, M. Systematic Evaluation of Methods for Integration of Transcriptomic Data into Constraint-Based Models of Metabolism. Plos Comput Biol 10 (2014). 10.1371/journal.pcbi.1003580

19. Zampieri, G., Vijayakumar, S., Yaneske, E. & Angione, C. Machine and deep learning meet genome-scale metabolic modeling. Plos Comput Biol 15 (2019). 10.1371/journal.pcbi.1007084

20. Kandalaft, L. E., Laniti, D. D. & Coukos, G. Immunobiology of high-grade serous ovarian cancer: lessons for clinical translation. Nat Rev Cancer 22, 640–656 (2022). 10.1038/s41568-022-00503-z

21. Bell, D. et al. Integrated genomic analyses of ovarian carcinoma. Nature 474, 609–615 (2011). 10.1038/nature10166

22. Bruchim, I. et al. Analyses of p53 expression pattern and BRCA mutations in patients with double primary breast and ovarian cancer. Int J Gynecol Cancer 14, 251–258 (2004). 10.1111/j.1048-891X.2004.014208.x

23. Ramus, S. J. & Gayther, S. A. The Contribution of BRCA1 and BRCA2 to Ovarian Cancer. Mol Oncol 3, 138–150 (2009). 10.1016/j.molonc.2009.02.001

24. Gudmundsdottir, K. & Ashworth, A. The roles of BRCA1 and BRCA2 and associated proteins in the maintenance of genomic stability. Oncogene 25, 5864–5874 (2006). 10.1038/sj.onc.1209874

25. Bruand, M. et al. Cell-autonomous inflammation of BRCA1-deficient ovarian cancers drives both tumor-intrinsic immunoreactivity and immune resistance via STING. Cell Rep 36 (2021). 10.1016/j.celrep.2021.109412

26. Bradbury, M. et al. BRCA1 mutations in high-grade serous ovarian cancer are associated with proteomic changes in DNA repair, splicing, transcription regulation and signaling. Sci Rep-Uk 12 (2022). 10.1038/s41598-022-08461-0

27. Werner, H. BRCA1: An Endocrine and Metabolic Regulator. Front Endocrinol (Lausanne) 13, 844575 (2022). 10.3389/fendo.2022.844575

28. Roig, B. et al. Metabolomics reveals novel blood plasma biomarkers associated to the BRCA1-mutated phenotype of human breast cancer. Sci Rep-Uk 7 (2017). 10.1038/s41598-017-17897-8

29. Cuyàs, E. et al. Germline mutation reprograms breast epithelial cell metabolism towards mitochondrial-dependent biosynthesis: Evidence for metformin-based “starvation” strategies in carriers. Oncotarget 7, 52974–52992 (2016). 10.18632/oncotarget.9732

30. Privat, M. et al. BRCA1 Induces Major Energetic Metabolism Reprogramming in Breast Cancer Cells. Plos One 9 (2014). 10.1371/journal.pone.0102438

31. Moreau, K. et al. BRCA1 affects lipid synthesis through its interaction with acetyl-CoA carboxylase. Journal of Biological Chemistry 281, 3172–3181 (2006). 10.1074/jbc.M504652200

32. Kanakkanthara, A. et al. BRCA1 Deficiency Upregulates NNMT, Which Reprograms Metabolism and Sensitizes Ovarian Cancer Cells to Mitochondrial Metabolic Targeting Agents. Cancer Res 79, 5920–5929 (2019). 10.1158/0008-5472.Can-19-1405

33. Lahiguera, A. et al. Tumors defective in homologous recombination rely on oxidative metabolism: relevance to treatments with PARP inhibitors. EMBO Mol Med 12, e11217 (2020). 10.15252/emmm.201911217

34. Bae, I. et al. BRCA1 induces antioxidant gene expression and resistance to oxidative stress. Cancer Res 64, 7893–7909 (2004). 10.1158/0008-5472.Can-04-1119

35. Neff, R. T., Senter, L. & Salani, R. BRCA mutation in ovarian cancer: testing, implications and treatment considerations. Ther Adv Med Oncol 9, 519–531 (2017). 10.1177/1758834017714993

36. Chiyoda, T. et al. Loss of BRCA1 in the Cells of Origin of Ovarian Cancer Induces Glycolysis: A Window of Opportunity for Ovarian Cancer Chemoprevention. Cancer Prev Res 10, 255–266 (2017). 10.1158/1940-6207.Capr-16-0281

37. Cucchi, D., Gibson, A. & Martin, S. A. The emerging relationship between metabolism and DNA repair. Cell Cycle 20, 943–959 (2021). 10.1080/15384101.2021.1912889

38. DelloRusso, C. et al. Functional characterization of a novel BRCA1-Null ovarian cancer cell line in response to ionizing radiation. Mol Cancer Res 5, 35–45 (2007). 10.1158/1541-7786.Mcr-06-0234

39. Schellenberger, J. et al. Quantitative prediction of cellular metabolism with constraint-based models: the COBRA Toolbox v2.0. Nat Protoc 6, 1290–1307 (2011). 10.1038/nprot.2011.308

40. Martinez-Outschoorn, U. E. et al. BRCA1 mutations drive oxidative stress and glycolysis in the tumor microenvironment Implications for breast cancer prevention with antioxidant therapies. Cell Cycle 11, 4402–4413 (2012). 10.4161/cc.22776

41. Argüello, R. J. et al. SCENITH: A Flow Cytometry-Based Method to Functionally Profile Energy Metabolism with Single-Cell Resolution. Cell Metabolism 32 (2020). 10.1016/j.cmet.2020.11.007

42. Nusinow, D. P. et al. Quantitative Proteomics of the Cancer Cell Line Encyclopedia. Cell 180, 387–402 e316 (2020). 10.1016/j.cell.2019.12.023

43. McCabe, A., Zaheed, O., McDade, S. S. & Dean, K. Investigating the suitability of in vitro cell lines as models for the major subtypes of epithelial ovarian cancer. Front Cell Dev Biol 11 (2023). 10.3389/fcell.2023.1104514

44. Pandey, V. & Hatzimanikatis, V. Investigating the deregulation of metabolic tasks via Minimum Network Enrichment Analysis (MiNEA) as applied to nonalcoholic fatty liver disease using mouse and human omics data. Plos Comput Biol 15 (2019). 10.1371/journal.pcbi.1006760

45. Vazquez-Garcia, I. et al. Ovarian cancer mutational processes drive site-specific immune evasion. Nature 612, 778–786 (2022). 10.1038/s41586-022-05496-1

46. Ruscito, I. et al. Characterisation of tumour microvessel density during progression of high-grade serous ovarian cancer: clinico-pathological impact (an OCTIPS Consortium study). Brit J Cancer 119, 330–338 (2018). 10.1038/s41416-018-0157-z

47. Ding, Y. et al. Molecular characteristics and tumorigenicity of ascites-derived tumor cells: mitochondrial oxidative phosphorylation as a novel therapy target in ovarian cancer. Mol Oncol 15, 3578–3595 (2021). 10.1002/1878-0261.13028

48. Bose, S. et al. G6PD inhibition sensitizes ovarian cancer cells to oxidative stress in the metastatic omental microenvironment. Cell Rep 39 (2022). 10.1016/j.celrep.2022.111012

49. Nieman, K. M. et al. Adipocytes promote ovarian cancer metastasis and provide energy for rapid tumor growth. Nat Med 17, 1498–U1207 (2011). 10.1038/nm.2492

50. Stine, Z. E., Schug, Z. T., Salvino, J. M. & Dang, C. V. Targeting cancer metabolism in the era of precision oncology. Nat Rev Drug Discov 21, 141–162 (2022). 10.1038/s41573-021-00339-6

51. Elia, I. & Haigis, M. C. Metabolites and the tumour microenvironment: from cellular mechanisms to systemic metabolism. Nat Metab 3, 21–32 (2021). 10.1038/s42255-020-00317-z

52. Stur, E. et al. The dynamic immune behavior of primary and metastatic ovarian carcinoma. NPJ Precis Oncol 9, 120 (2025). 10.1038/s41698-025-00818-8

53. Mikulak, J. et al. Immune evasion mechanisms in early-stage I high-grade serous ovarian carcinoma: insights into regulatory T cell dynamics. Cell Death Dis 16 (2025). 10.1038/s41419-025-07557-5

54. Tsai, C. H. et al. Immunoediting instructs tumor metabolic reprogramming to support immune evasion. Cell Metabolism 35, 118-+ (2023). 10.1016/j.cmet.2022.12.003

55. Chang, C. H. et al. Metabolic Competition in the Tumor Microenvironment Is a Driver of Cancer Progression. Cell 162, 1229–1241 (2015). 10.1016/j.cell.2015.08.016

56. Ritchie, M. E. et al. limma powers differential expression analyses for RNA-sequencing and microarray studies. Nucleic Acids Res 43, e47 (2015). 10.1093/nar/gkv007

57. Hänzelmann, S., Castelo, R. & Guinney, J. GSVA: gene set variation analysis for microarray and RNA-Seq data. Bmc Bioinformatics 14 (2013). 10.1186/1471-2105-14-7

58. Salvy, P. et al. pyTFA and matTFA: a Python package and a Matlab toolbox for Thermodynamics-based Flux Analysis. Bioinformatics 35, 167–169 (2019). 10.1093/bioinformatics/bty499

59. Gudmundsson, S. & Thiele, I. Computationally efficient flux variability analysis. Bmc Bioinformatics 11 (2010). 10.1186/1471-2105-11-489

60. Heirendt, L. et al. Creation and analysis of biochemical constraint-based models using the COBRA Toolbox v.3.0. Nat Protoc 14, 639–702 (2019). 10.1038/s41596-018-0098-2

61. Bedia, J. S. et al. Estimating the ovarian cancer CA-125 preclinical detectable phase, in-vivo tumour doubling time, and window for detection in early stage: an exploratory analysis of UKCTOCS. Ebiomedicine 112 (2025). 10.1016/j.ebiom.2024.105554

62. van der Velpen, V. et al. Systemic and central nervous system metabolic alterations in Alzheimer’s disease. Alzheimers Res Ther 11 (2019). 10.1186/s13195-019-0551-7

63. Gallart-Ayala, H. et al. A global HILIC-MS approach to measure polar human cerebrospinal fluid metabolome: Exploring gender-associated variation in a cohort of elderly cognitively healthy subjects. Anal Chim Acta 1037, 327–337 (2018). 10.1016/j.aca.2018.04.002

64. Broadhurst, D. et al. Guidelines and considerations for the use of system suitability and quality control samples in mass spectrometry assays applied in untargeted clinical metabolomic studies. Metabolomics 14, 72 (2018). 10.1007/s11306-018-1367-3

