## Supplementary Figures for "Computational Inference of Metabolic Programs: A Case Study Analyzing the Effect of BRCA1 Loss"

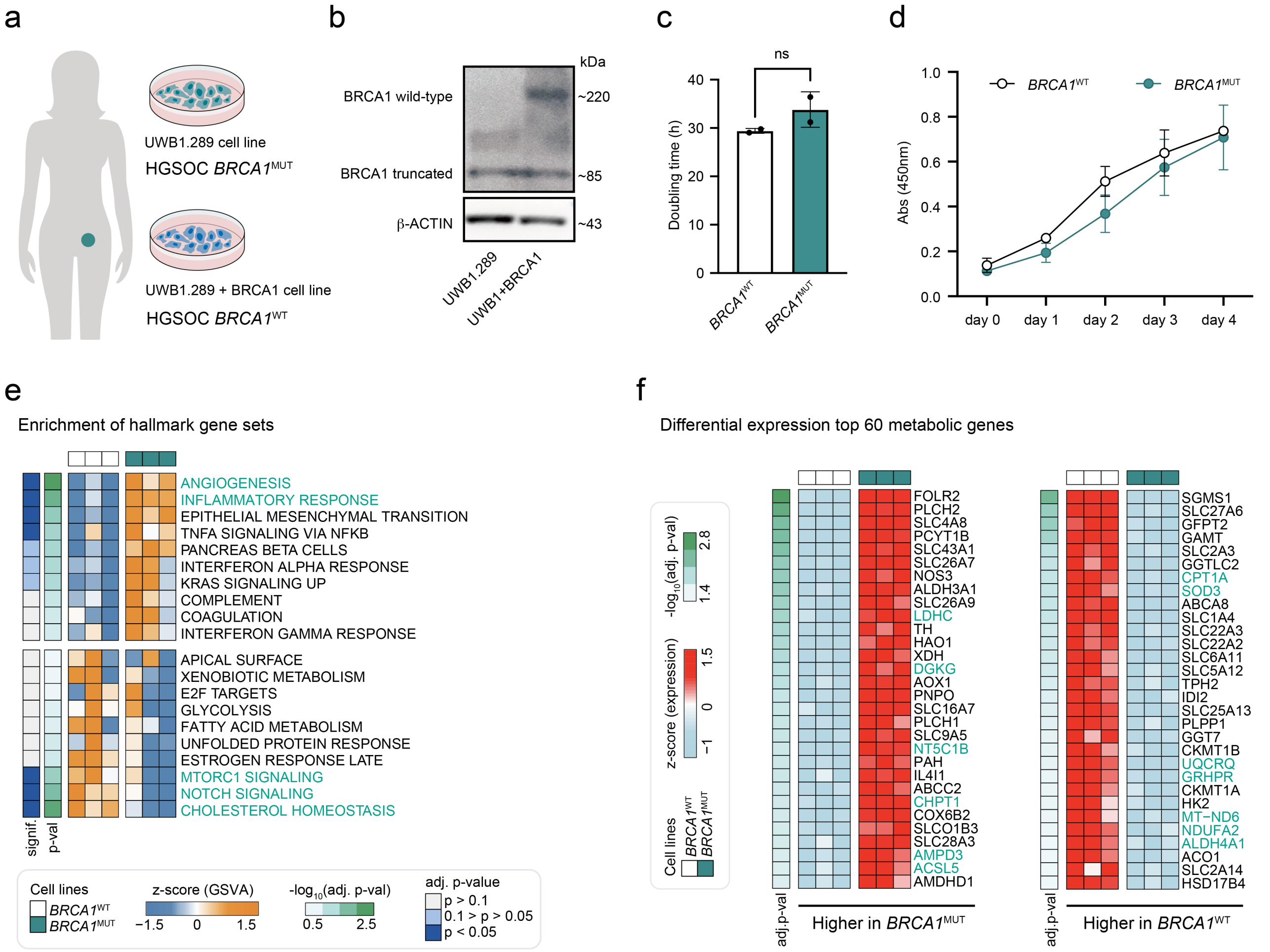


**Figure S1. BRCA1 loss alters metabolic gene expression in HGSOC UWB1.289 cell line. a,** BRCA1-deficient (UWB1.289) and wild-type BRCA1-restored (UWB1.289+BRCA1) HGSOC cell lines. **b,** Wild-type and truncated BRCA1 protein expression in the UWB1.289 and UWB1.289+BRCA1 cell lines. **c,** Doubling time of the UWB1.289+BRCA1 (*BRCA1*$^{WT}$) and UWB1.289 (*BRCA1*$^{MUT}$) cell lines (n=2 each in triplicate). **d,** Growth assay of the *BRCA1*$^{WT}$ and *BRCA1*$^{MUT}$ UWB1.289 cell lines (n=3). **e,** Hallmarks signatures (as defined in MSigDB) enrichment analysis calculated applying GSVA with bulk RNA-seq data for *BRCA1*$^{MUT}$ and *BRCA1*$^{WT}$ cell lines (n=3). **f,** Top 60 significantly (p<0.05) differentially expressed metabolic genes between *BRCA1*$^{MUT}$ and *BRCA1*$^{WT}$ cell lines (n=3). Statistical comparisons were calculated by unpaired t-test FDR adjusted.

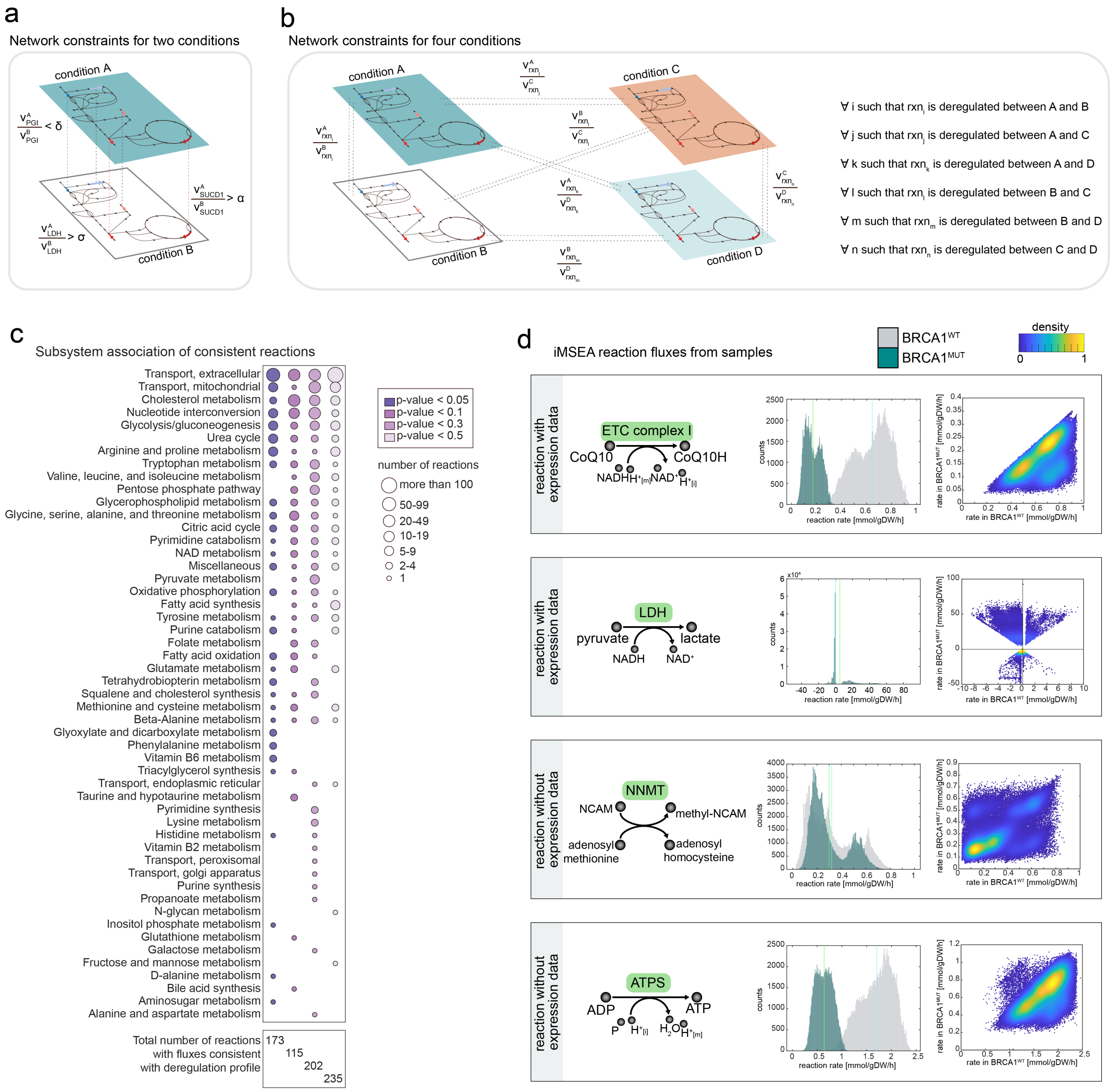


**Figure S2. iMSEA data integration and metabolic flux sampling. a-b,** Scheme of network constraints (as used in REMI and iMSEA) to integrate pairwise deregulated ratios for two conditions (**a**) or for four conditions (**b**). **c,** Number of reactions with fluxes consistent with expression profile classified by metabolic subsystem and integration step. **d,** Examples of iMSEA flux samples for reactions with expresion data (reactions constrained according to deregulation profile) and for reactions without expression data (fluxes inferred from the network).

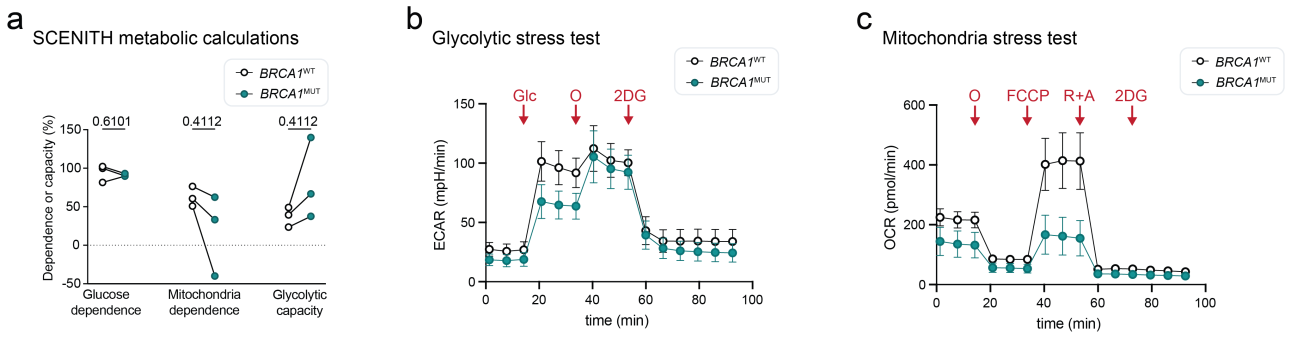


**Figure S3. Experimental validation of metabolic state in UWB1.289 cell lines. a,** Calculations of metabolic profile of *BRCA1*$^{WT}$ and *BRCA1*$^{MUT}$ cells after SCENITH analysis. Percentage of dependence or capacity is shown (n=3 independent experiments, each in 3-6 replicates). **b,** ECAR assessment in *BRCA1*$^{WT}$ and *BRCA1*$^{MUT}$ cells by glycolytic stress test experiment (n=10). **c,** OCR assessment in *BRCA1*$^{WT}$ and *BRCA1*$^{MUT}$ cells by mitochondrial stress test seahorse experiment (n=10).

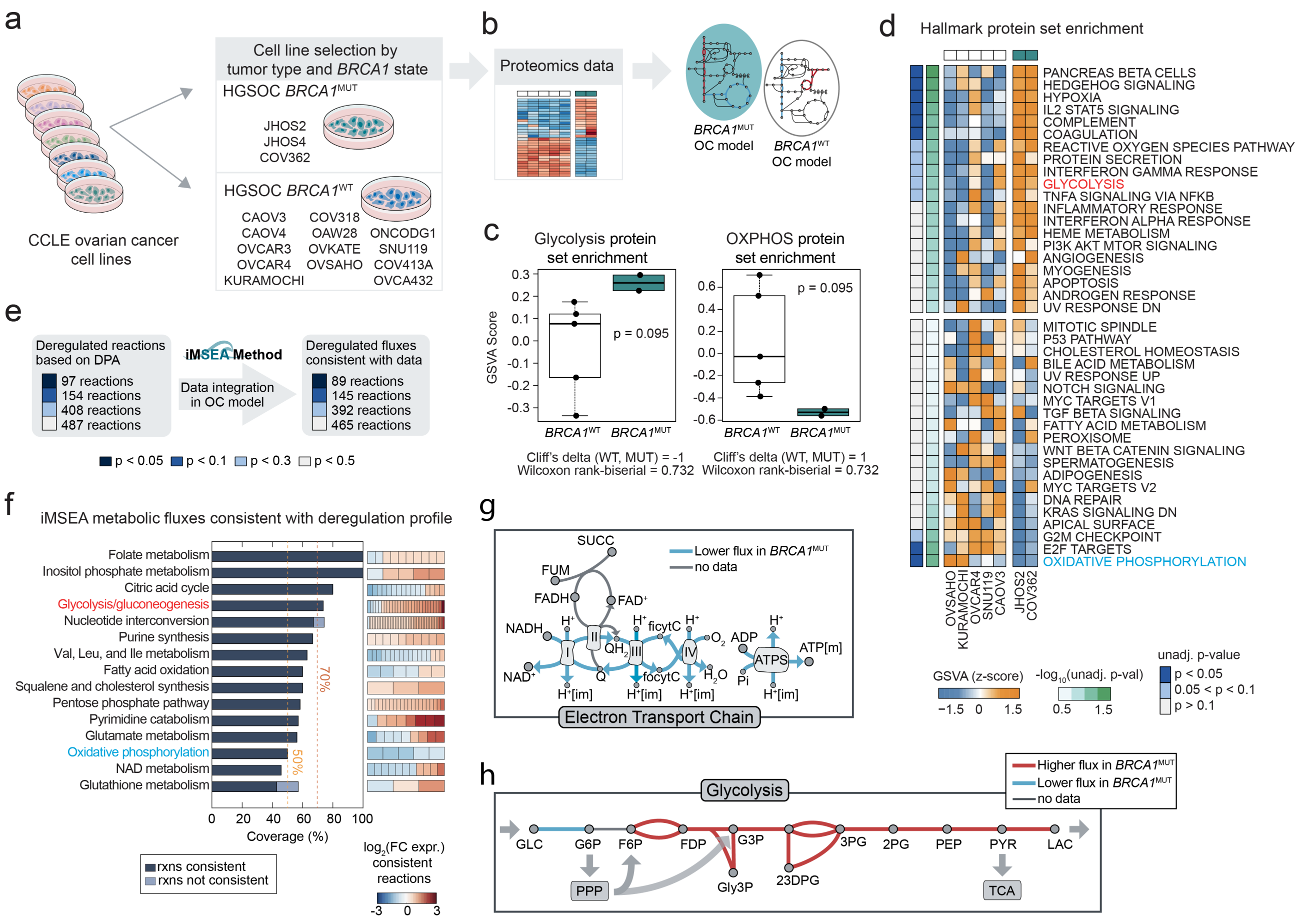


**Figure S4. Metabolic states across BRCA1-deficient and WT ovarian cancer cell lines from the CCLE. a,** Cell line selection based on tumor type and *BRCA1* status. **b,** Integration of differential protein abundance for *BRCA1*$^{WT}$ (n=5) and *BRCA1*$^{MUT}$ (n=2) cells into ovarian cancer metabolic network. **c,** Enrichment score of glycolysis and oxidative phosphorylation (OXPHOS) pathway calculated by applying GSVA with available proteomics data of *BRCA1*$^{WT}$ and *BRCA1*$^{MUT}$ cells. **d,** Hallmarks gene set enrichment analysis calculated applying GSVA to protein abundances. **e,** Mapping deregulated genes (proteomics) to reactions in the ovarian cancer model, by using gene-protein-reaction rules. **f,** Metabolic subsystems with reaction fluxes consistent with expression profile, including coverage of subsystem and fold change expression of reactions in BRCA1$^{MUT}$ vs. BRCA1$^{WT}$ cell lines. **g-h,** Network representation of **g,** the electron transport chain and **h,** glycolysis with consistent (based on proteomics data) downregulated reactions colored in blue, upregulated reactions colored in red, and reactions lacking data colored in grey. Statistical comparisons calculated by Wilcoxon Rank-Sum test (**c**), or unpaired t test FDR adjusted (**d**).

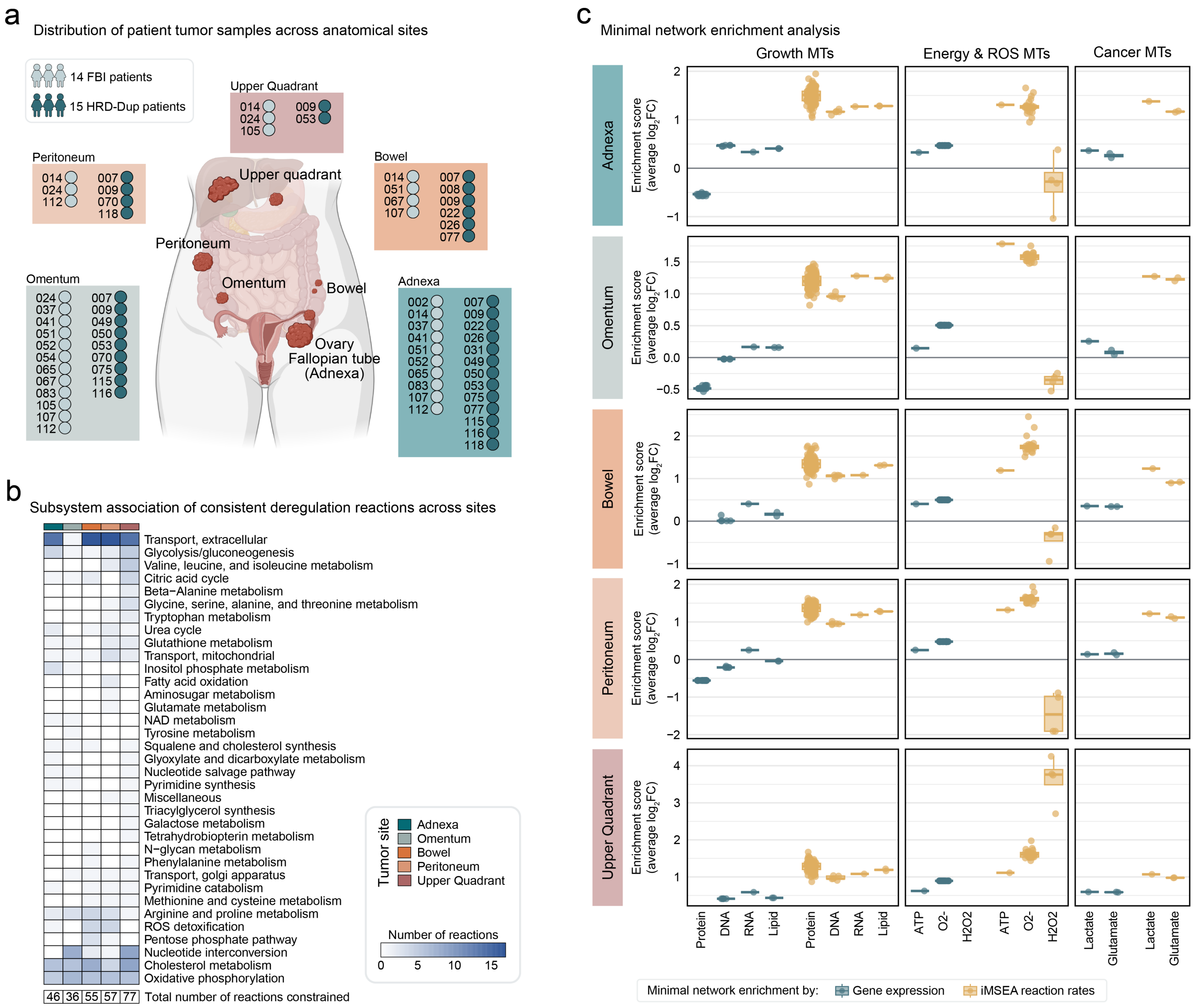


**Figure S5. Analysis of metabolic states for HRD-Dup and FBI patients across sites. a,** Distribution of samples from the cohort of HGSOC patients by site and homologous recombination status. **b,** Number of deregulated reactions (in HRD-Dup vs. FBI patients) with fluxes consistent with the expression profile classified by metabolic subsystem across sites. **c,** Minimal network enrichment analysis of ovarian cancer metabolic tasks across tumor sites using gene expression data or iMSEA reaction rates.

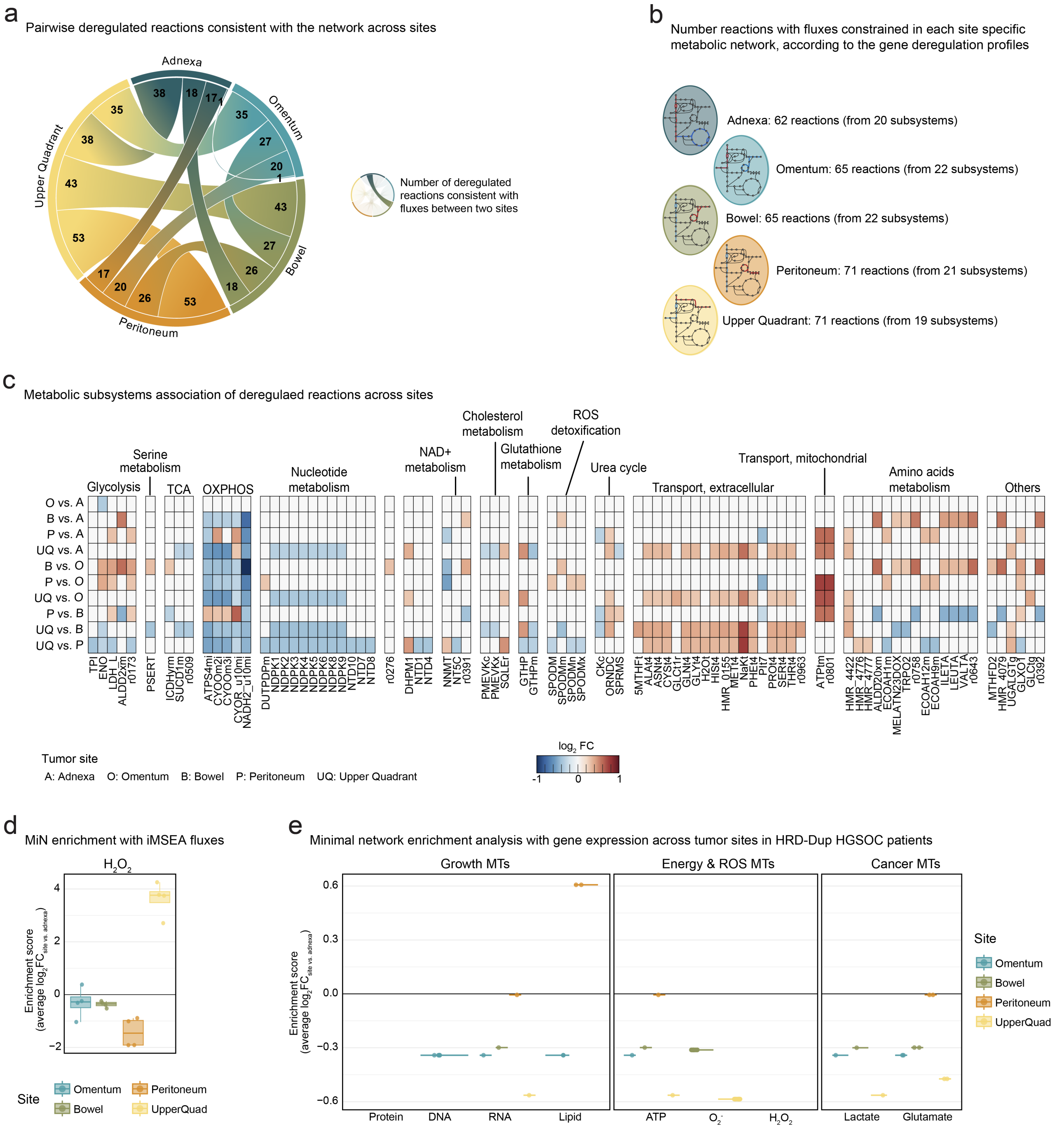


**Figure S6. Metabolic landscapes across tumor sites in HRD-Dup HGSOC patients. a,** Number of pairwise deregulated reactions consistent with the network across anatomical sites of HRD-Dup patients. **b,** Number of reactions with fluxes constrained by the gene deregulation profile in each site-specific metabolic network. **c,** Association to metabolic subsystems of reactions constrained by gene expression in site-specific networks, showing $\log_2$ fold changes (FC) in reaction rates constrained by upregulated (red) and downregulated (blue) genes when comparing gene expression pairwise across different sites. **d,** Minimal network enrichment with iMSEA fluxes of the hydrogen peroxide metabolic task across sites for HRD-Dup patients. **e,** Minimal network enrichment analysis of metabolic tasks with gene expression data across tumor sites for HRD-Dup patients.
